# Stabilizing genetically unstable simple sequence repeats in the *Campylobacter jejuni* genome by multiplex genome editing: a reliable approach for delineating multiple phase-variable genes

**DOI:** 10.1101/2021.05.14.444138

**Authors:** Shouji Yamamoto, Sunao Iyoda, Makoto Ohnishi

## Abstract

Hypermutability of simple sequence repeats (SSR) through DNA slippage is a major mechanism of phase variation in *Campylobacter jejuni.* The presence of multiple SSR-mediated phase-variable genes encoding enzymes that modify surface structures, including capsular polysaccharide (CPS) and lipooligosaccharide (LOS), generates high levels of structural variants within bacterial populations, thereby promoting adaptation to selective pressures in host environments. Therefore, the phenotypic diversity generated by phase variation can limit the reproducibility of results with *C. jejuni;* therefore, researchers need to genetically control the mutability of multiple SSRs. Here, we show that natural “cotransformation” is an effective method for *C. jejuni* genome editing. Cotransformation is a trait of naturally competent bacteria that causes uptake and integration of multiple different DNA fragments, which has been recently adapted to multiplex genome editing by natural transformation (MuGENT), a method for introducing multiple scarless mutations into the genomes of these bacteria. We found that the cotransformation frequencies of antibiotic resistance gene-marked DNA fragments and unmarked DNA fragments reached ~40% in *C. jejuni.* To examine the feasibility of MuGENT in *C. jejuni,* we “locked” either different polyG SSR tracts in strain NCTC11168 (which are located in the biosynthetic CPS and LOS gene clusters) into either the ON or OFF configurations by interrupting the continuous runs of G residues without changing the encoded amino acids. This approach, termed “MuGENT-SSR,” enabled the generation of all eight edits within 2 weeks and the identification of a phase-locked strain with a highly stable type of Penner serotyping, a CPS-based serotyping scheme. Furthermore, extensive genome editing of this strain by MuGENT-SSR identified a phase-variable gene that determines the Penner serotype of NCTC11168. Thus, MuGENT-SSR provides a platform for genetic and phenotypic engineering of genetically unstable *C. jejuni,* making it a reliable approach for elucidating the mechanisms underlying phase-variable expression of specific phenotypes.

**Author summary:** *Campylobacter jejuni* is the leading bacterial cause of food-borne gastroenteritis in developed countries and occasionally progresses to the autoimmune disease Guillain–Barré syndrome. The genetically and phenotypically unstable features of this bacterial species limit research and development efforts. A relatively large number of hypermutable simple sequence repeat (SSR) tracts in the *C. jejuni* genome markedly decreases its phenotypic stability through reversible changes in the ON or OFF expression states of the genes in which they reside, a phenomenon called phase variation. Thus, controlling SSR-mediated phase variation can be important for achieving stable and reproducible research on *C. jejuni*. In this study, we developed a feasible and effective approach to genetically manipulate multiple SSR tracts in the *C. jejuni* genome using natural cotransformation, a trait of naturally transformable bacterial species that causes the uptake and integration of multiple different DNA molecules. This approach will greatly help to improve the genetic and phenotypic stability of *C. jejuni* to enable diverse applications in research and development.

## Introduction

Simple sequence repeats (SSR) or microsatellites in bacterial genomes are highly mutable because of the potential for DNA-strand slippage during DNA replication [1]. Mispairing DNA by slippage results in the insertion or deletion of one or more repeats within a repetitive DNA element. When an SSR tract is present within an open reading frame (ORF), promoter, or other regulatory sequences, its hypermutability mediates reversible and frequent changes in specific phenotypes through transcriptional or translational genetic switches between ON and OFF states, which is called “phase variation” [2,3]. In several bacterial species, including *Campylobacter jejuni,* the presence of multiple SSR-mediated phase-variable genes (also called contingency genes) per genome generates high levels of phenotypic variants within bacterial populations [4,5].

*C. jejuni* is the leading bacterial cause of food-borne gastroenteritis in developed countries, primarily depending on its ability to colonize the caeca of chickens and to survive in the food chain by attaching to undercooked chicken meat [6,7]. *C. jejuni* readily colonizes the intestinal mucosa of a wide variety of wild and domestic birds and other animals. Infections in poultry are usually asymptomatic, while human infection can cause significant inflammation and bloody diarrhea, occasionally progressing to the autoimmune disease, Guillain–Barré syndrome [8]. As a commensal bacterium of poultry and a human pathogen, *C. jejuni* needs to rapidly adapt to differences in host environments, such as changing nutrient compositions and immune systems. Additional selective pressures are caused by transmission through genetically and immunologically variable host populations and exposure to bacteriophages. *C. jejuni* likely utilizes phase variation as a major mechanism for adapting to these selective pressures.

A relatively large number of SSR tracts consisting of seven or more G or C bases was unexpectedly found in the AT-rich genomes of *C. jejuni.* These polyG/C SSR tracts are mainly involved in phase variation in this species, and a genomic analysis of four *C. jejuni* strains indicated the presence of 12 to 29 tracts per genome [9]. *C. jejuni* NCTC11168 encodes 29 polyG/C tracts, 23 of which are in protein-coding regions [10], suggesting that phase variation in this strain is regulated mainly at the translational level. The majority of these loci are clustered on the genomic regions predicted to encode enzymes involved in modifying surface structures, including lipooligosaccharide (LOS), capsular polysaccharide (CPS), and flagella, but a few are located on genes encoding cell-surface proteins or restriction enzymes [5]. The ability to change the antigenicity of surface structures by phase variation strongly suggests that this phenotypic diversity could affect colonization or pathogenesis in host organisms. Consistently, it has been reported that phase-variable expression of specific genes affects pathogenesis in humans and colonization in chickens [11–15]. Comprehensive studies of multiple phase-variable genes in *C. jejuni* have also demonstrated that culturing or passage through animal and human hosts results in significant phase changes in multiple SSR-containing genes, particularly those involved in surface-structure modifications, which in some cases, are coupled with enhanced colonization and pathogenesis [9,16–20].

A limitation of studying the contributions of multiple SSR-mediated phase-variable genes to specific phenotypes is that researchers currently lack genetic tools for efficiently “locking” SSR into either ON or OFF states at multiple loci in the *C. jejuni* genome. For example, the use of a multiplexed SSR-editing technology enables phenotype stabilization and therefore promotes reproducible research activities in this area. Such phenotypic engineering would also be applicable for stably producing phase-variable surface antigenic determinants, which can be used for vaccine development and raising serotyping antisera. To date, multiplex automated genome engineering (MAGE) using oligonucleotides, a highly efficient λRed recombinase, and clustered regularly interspaced short palindromic repeats (CRISPR)/CRISPR-associated protein (Cas) systems have been developed for targeted genome engineering in bacteria [21,22]. However, these methods require not only engineered plasmids or chromosomes for the respective genome-editing systems (i.e., expressing proteins and/or RNAs involved in the λRed and CRISPR/Cas systems) but also, in the case of CRISPR/Cas, the selection of edited genomic sites. Natural competence for transformation is shared by diverse bacterial species [23,24]. This process involves the uptake of exogenous DNA, followed by integration of the imported DNA into the genome by homologous recombination. Because natural transformation does not require any special factors and is dependent on the recipient bacterium and donor DNA molecules, it has long been used for genetic engineering in naturally competent bacterial species. *C. jejuni* was shown to be naturally transformable over 30 years ago [25]. However, natural transformation has not been conventionally used as a genetic engineering method in this bacterium because it can be highly transformed with DNA prepared from *C. jejuni*, but not with DNA propagated in *Escherichia coli* or amplified by the polymerase chain reaction (PCR). Recently, Beauchamp *et al*. demonstrated that methylation at the RAATTY sequence of *E. coli*-derived plasmid DNA or PCR-amplified DNA can efficiently transform *C. jejuni* [26]. In a study of other transformable species conducted by Dalia *et al.,* the authors established the multiplex genome editing by natural transformation (MuGENT) system, which is based on “cotransformation,” a trait that causes the uptake and integration of multiple different DNA molecules [27–29]. These two pioneering studies have provided great potential for performing multiplexed gene modifications in *C. jejuni* and motivated us to examine this feasibility. Here, we optimized natural transformation and cotransformation using PCR-amplified donor DNA fragments and demonstrated its utility as a method for multiplex genome editing in naturally competent *C. jejuni.*

## Results and discussion

### Optimization of donor DNA for the natural transformation of *C. jejuni*

Data from a previous study by Beauchamp *et al.* demonstrated that DNA methylated at the RA^m6^ATTY sequence is an efficient substrate for *C. jejuni* transformation [26]. To optimize the natural transformation of this bacterium, we investigated in greater detail the characteristics of DNA substrate generated using PCR products. For this purpose, we amplified various DNA fragments of the *rpsL*^K88R^ allele, which confers streptomycin resistance (Sm^R^) [30]. These *rpsL*^K88R^ marker fragments had different lengths of regions homologous to the recombination target sequence (50–2,000 base pairs [bp]) and different numbers of the EcoRI methyltransferase-recognition sequence (GAATTC), which was endogenously present or exogenously added to the PCR primers (Fig 1, n = 1 to 3). Each 0.5 pmol of the amplified DNA was treated with EcoRI methyltransferase and then used in transformation assays to transform the NCTC11168 and 81-176 strains, which are representative strains used for genetic studies of *C. jejuni* [5,31,32] (Table 1). We did not obtain transformants using fragments with short regions of homology (50 or 100 bp), even though these DNA fragments had two methylated GAATTC sites (Fig 1, *rpsL*^K88R^-1 and *rpsL*^K88R^-2). In contrast, the NCTC11168 and 81-176 strains were transformed at substantially higher frequencies using fragments with longer regions of homology (500 bp; Fig 1, *rpsL*^K88R^-5). The highest transformation frequencies were obtained when 1,000-to 2,000-bp homologies were present (Fig 1, *rpsL*^K88R^-8 and *rpsL*^K88R^-9). The length of a homologous sequence is a major determinant of the efficiency of RecA-dependent recombination [33]. In the *ΔrecA* background, no transformants were detected using a fragment with 2,000 homologous bp (Fig 1, *rpsL*^K88R^-9), suggesting that transformation in *C. jejuni* is mediated by RecA. We also confirmed that the lack of the GAATTC sequence or the lack of GAATTC methylation markedly decreased the transformation frequency (Fig 1, *rpsL*^K88R^-3 and unmethylated *rpsL*^K88R^-5). However, increasing the number of methylated GAATTC sites did not substantially increase the transformation frequencies (Fig 1, *rpsL*^K88R^-4 and *rpsL^K88R^-5* or *rpsL^K88R^-6, rpsL^K88R^-7,* and *rpsL^K88R^-8).* These results suggest that one methylated GAATTC sequence is sufficient for transformation, consistent with previous results obtained using plasmid DNA as a donor [26]. In summary, to maximize the efficiency of *C. jejuni* transformation, the donor DNA should contain two key structural elements: (1) a ≥1,000-bp region of homology and (2) at least one methylated GAATTC site.

**Fig 1.**
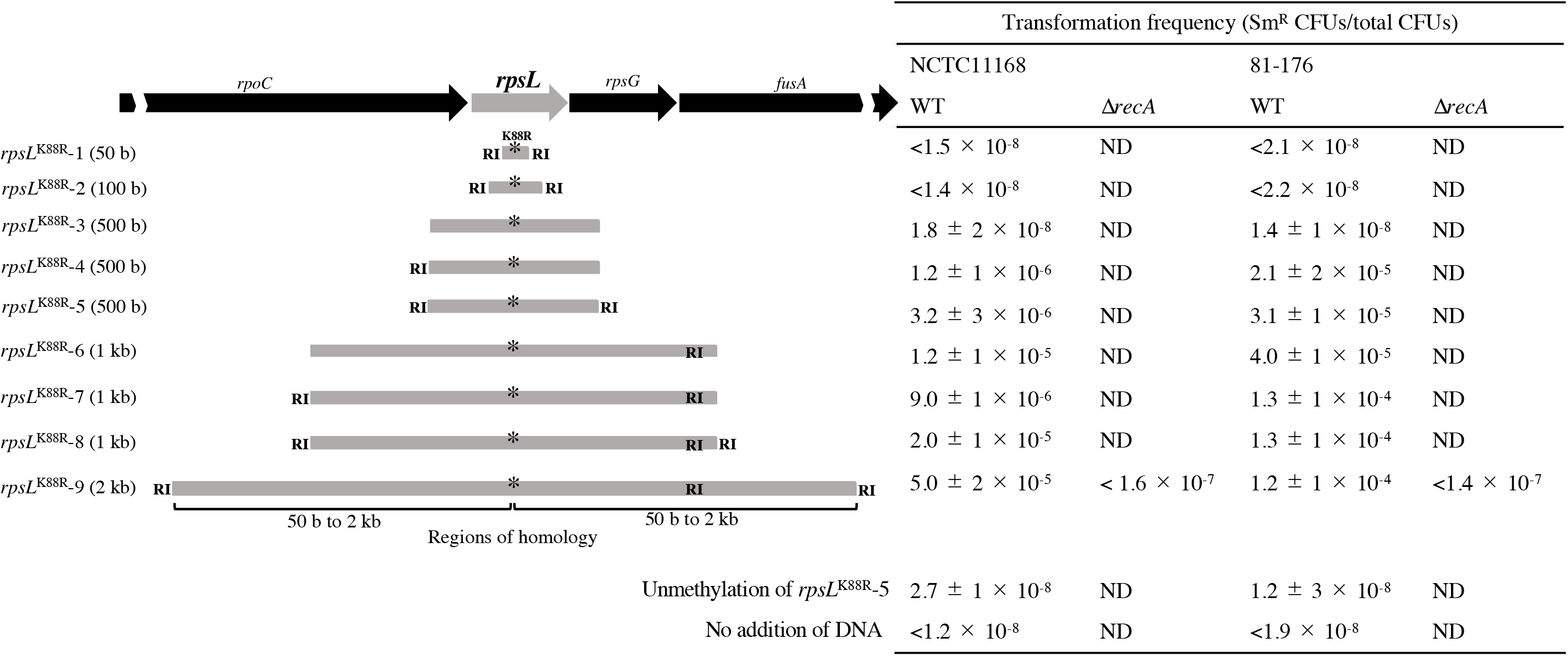
Effects of homology length and methylation on natural transformation. *C. jejuni* cells were transformed using *rpsL*^K88R^ marker fragments with different lengths of homologous regions compared to the recombination target sequence (50–2,000 bp, indicated in parentheses) and different numbers of EcoRI methyltransferase-recognition sequences (RIs). Each 0.5 pmol of amplified DNA was treated with EcoRI methyltransferase and then used in the transformation assays. The transformation frequency was defined as the number of Sm^R^ colony-forming units (CFUs) divided by the total number of CFUs. Data from three independent transformations are presented as the means ± standard deviations. As controls, we used unmethylated DNA (indicated by “Unmethylation of *rpsL*^K88R^-5”) or assayed for transformation in the absence of donor DNA (indicated by “No addition of DNA”). “ND” indicates not done. The following strains were used: NCTC11168 WT, NCTC11168; NCTC11168 *ΔrecA,* SYC1004; 81-176 WT, 81-176; 81-176 Δ*recA*, and SYC2004. The following DNA fragments were used (S2 Table): *rpsL*^K88R^-1 and *rpsL*^K88R^-2 (for both strains); *rpsL*^K88R^-3 and *rpsL*^K88R^-3-1 (for NCTC11168) and *rpsL*^K88R^-3-2 (for 81-176); *rpsL*^K88R^-4 and *rpsL*^K88R^-4-1 (for NCTC11168) and *rpsL*^K88R^-4-2 (for 81-176); *rpsL*^K88R^-5 and *rpsL*^K88R^-5-1 (for NCTC11168) and *rpsL*^K88R^-5-2 (for 81-176); *rpsL*^K88R^-6 and *rpsL*^K88R^-6-1 (for NCTC11168) and *rpsL*^K88R^-6-2 (for 81-176); *rpsL*^K88R^-7 and *rpsL*^K88R^-7-1 (for NCTC11168) and *rpsL*^K88R^-7-2 (for 81-176); *rpsL*^K88R^-8 and *rpsL*^K88R^-8-1 (for NCTC11168) and *rpsL*^K88R^-8-2 (for 81-176); and *rpsL*^K88R^-9 and *rpsL*^K88R^-9-1 (for NCTC11168) and *rpsL*^K88R^-9-2 (for 81-176).

**Table 1.**
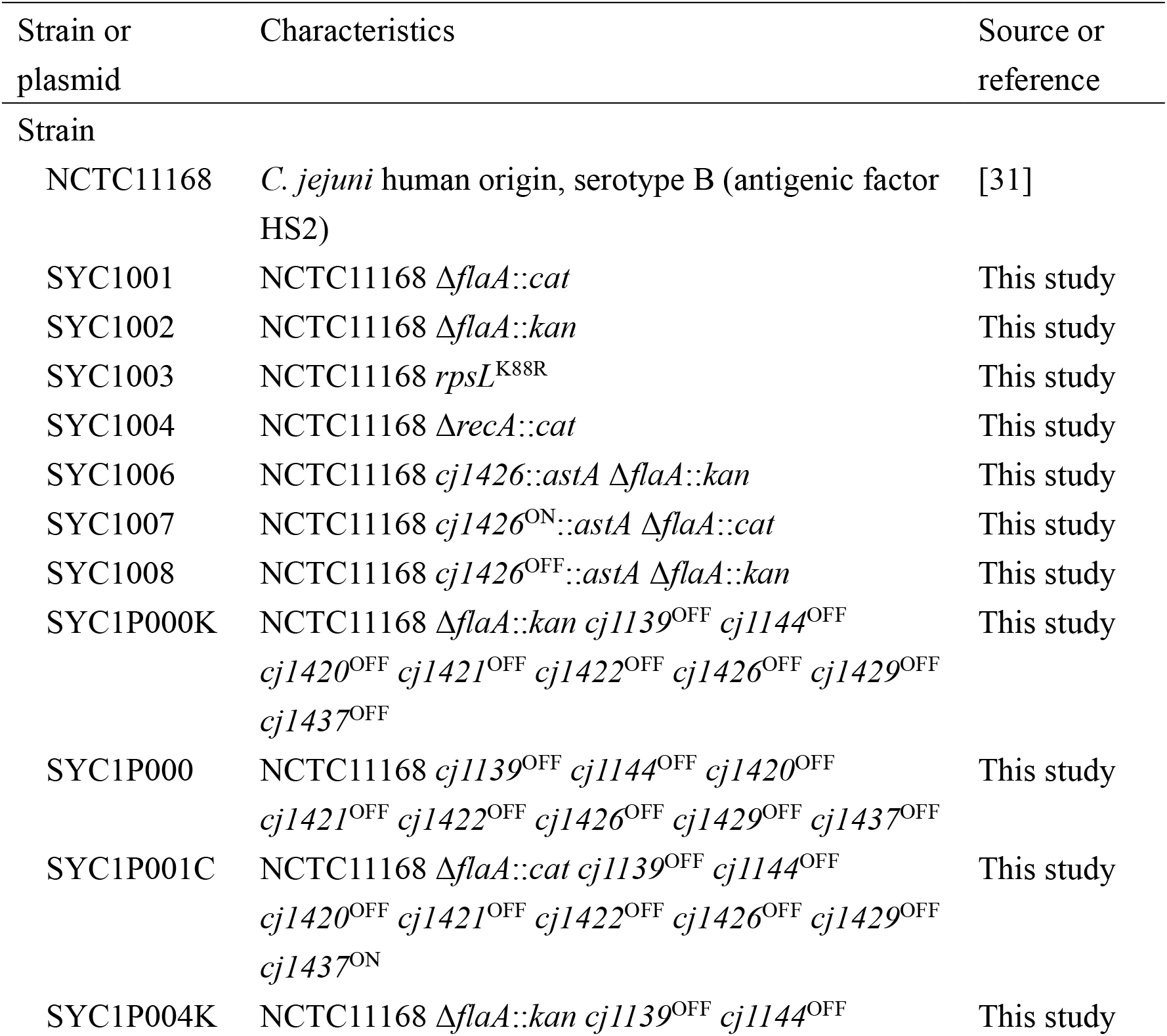
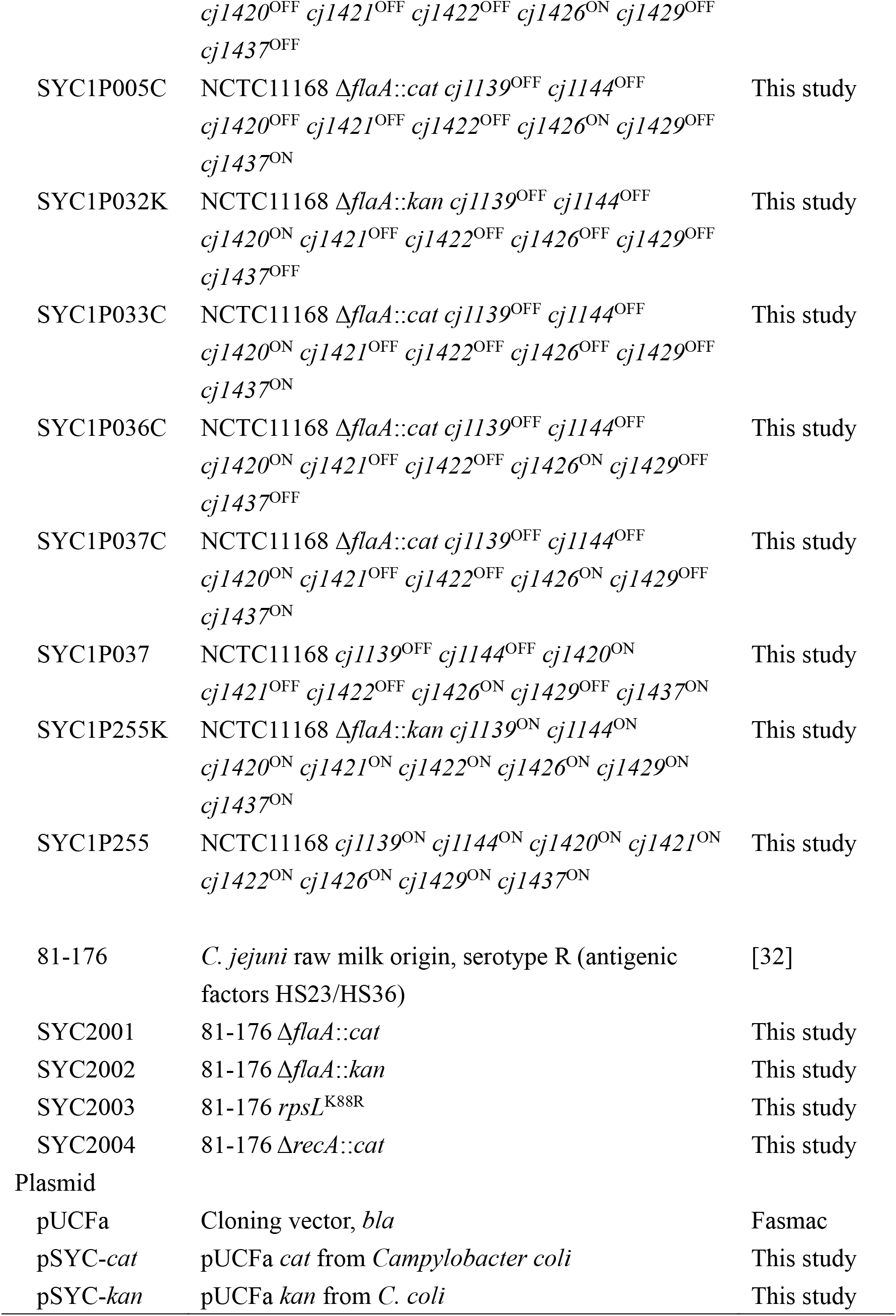
Strains and plasmids used in this study.

### Occurrence of natural cotransformation in *C. jejuni*

Using *Vibrio cholerae* and *Streptococcus pneumoniae,* Dalia *et al.* developed the MuGENT system using cotransformation, a trait of several naturally competent species [27–29]. During MuGENT, a bacterial culture is incubated with two types of donor DNA fragments: (1) a selected fragment that introduces an antibiotic-resistance gene into the genome and (2) unselected fragments that introduce scarless or transgene-free edits of interest at one or more loci. In *V. cholerae,* the frequencies of cotransformation of these distinct genetic markers can be made to exceed 60% by increasing the length of homology and the concentration of the unselected fragment [27]. To assess natural cotransformation in *C. jejuni,* we used two strains (SYC1003 and SYC2003) as recipients, which harbor the *rpsL*^K88R^ mutations in the NCTC11168 and 81-176 genomes, respectively, and therefore are resistant to Sm (Fig 2). We also used a *ΔflaA::kan* fragment with 1,000-bp regions of homology to replace the flagellin gene with a kanamycin-resistance (Km^R^) marker (selected) and *rpsL^+^* fragments with homologous regions of different sizes (1,000 bp: *rpsL*^+^-8, 2,000 b: *rpsL*^+^-9) to revert to the wild-type Sm-sensitive (Sm^S^) phenotype (unselected). After transforming SYC1003 and SYC2003 with equimolar concentrations of the selected and unselected methylated DNA (mDNA) fragments, we selected Km^R^-transformants, 100 of which were then subjected to Sm-sensitivity testing to evaluate cotransformation (Fig 2). Using unselected mDNA with a 1,000-bp region of homology resulted in cotransformation frequencies of 24% and 16% in SYC1003 and SYC2003, respectively, and acquisition of the Km^R^ and Sm^S^ phenotypes (Fig 2). Furthermore, increasing the homology of unselected mDNA (2,000 bp) slightly increased the cotransformation frequencies of both strains (Fig 2). Thus, we recommend the use of unselected mDNA with 2,000-bp homology for efficient *C. jejuni* cotransformation.

**Fig 2.**
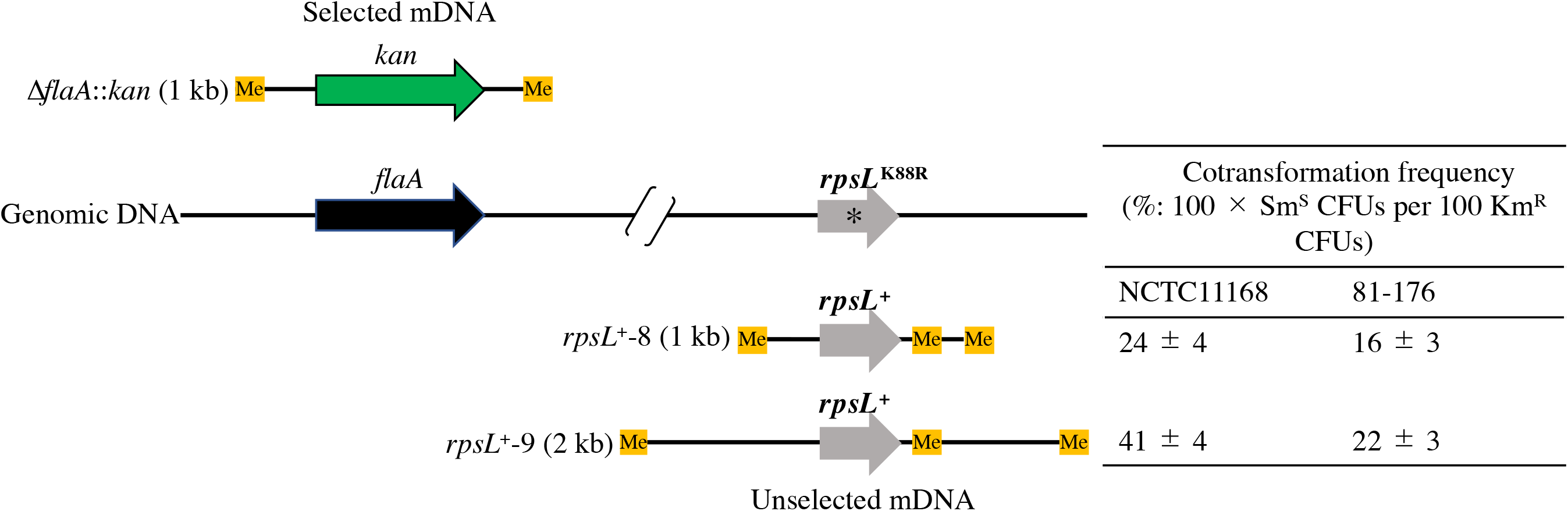
Evaluation of natural *C. jejuni* cotransformation. Natural cotransformation of *C. jejuni* was examined using two different PCR fragments, namely Δ*flaA::kan* and *rpsL*^+^ (S2 Table). The former fragments (Δ*flaA::kan-1* for NCTC11168, or Δ*flaA::kan-2* for 81-176) had 1,000 bp regions of homology (selected), whereas the latter fragments (*rpsL*^+^-8-1 and *rpsL*^+^-9-1 for NCTC11168, or *rpsL*^+^-8-2 and *rpsL*^+^-9-2 for 81-176) had 1,000-b or 2,000-b regions of homology (unselected). “Me” indicates a methylated site. Equimolar quantities (0.5 pmol) of selected and unselected fragments were independently methylated (indicated by the labeling “Selected mDNA” and “Unselected mDNA,” respectively), mixed, and then purified. After transforming strains carrying the *rpsL*^K88R^ allele (SY1003 and SY2003) with these mDNA fragments, the bacteria were spread onto brain heart infusion (BHI) agar plates containing Km. One hundred Km^R^-transformants were replica-plated onto BHI agar plates containing Sm. The cotransformation frequency (%) was calculated as follows: 100 × number of Sm^S^ CFUs per 100 Km^R^ CFUs. Data from three independent transformations are presented as the means ± standard deviations.

Natural cotransformation is thought to reflect the nature of competent bacterial cells, among which only a subpopulation of cells in a culture is transformable [28]. We demonstrated for the first time that *C. jejuni* cells are capable of cotransformation. It will be important to elucidate whether *C. jejuni* cells show variable competence within a population.

### Establishment of MuGENT in *C. jejuni*

MuGENT provides methods for simultaneously generating multiple scarless mutations and can therefore be broadly applied in diverse research and biotechnology applications [27]. We wanted to test whether natural cotransformation could be used for multiplex genome editing of *C. jejuni.* Fig 3 presents a schematic representation of our strategy. Briefly, multiple unselected mDNA fragments (used for introducing genome edits of interest) were mixed at equimolar concentrations with a selected mDNA fragment, and the resulting mixture was used to cotransform *C. jejuni* cells. Because MuGENT often requires multiple cycles of cotransformation in order to complete the genome editing, different selectable markers should be used for each cycle [27]. We swapped different resistance markers at the *flaA* gene at every MuGENT cycle. This also enabled easy removal of the marker genes and reversion to the wild-type allele by transformation with *flaA^+^* mDNA and subsequent selection of motile clones. Genome editing was verified by multiplex allele-specific colony (MASC) PCR [34] and nucleotide sequencing.

**Fig 3.**
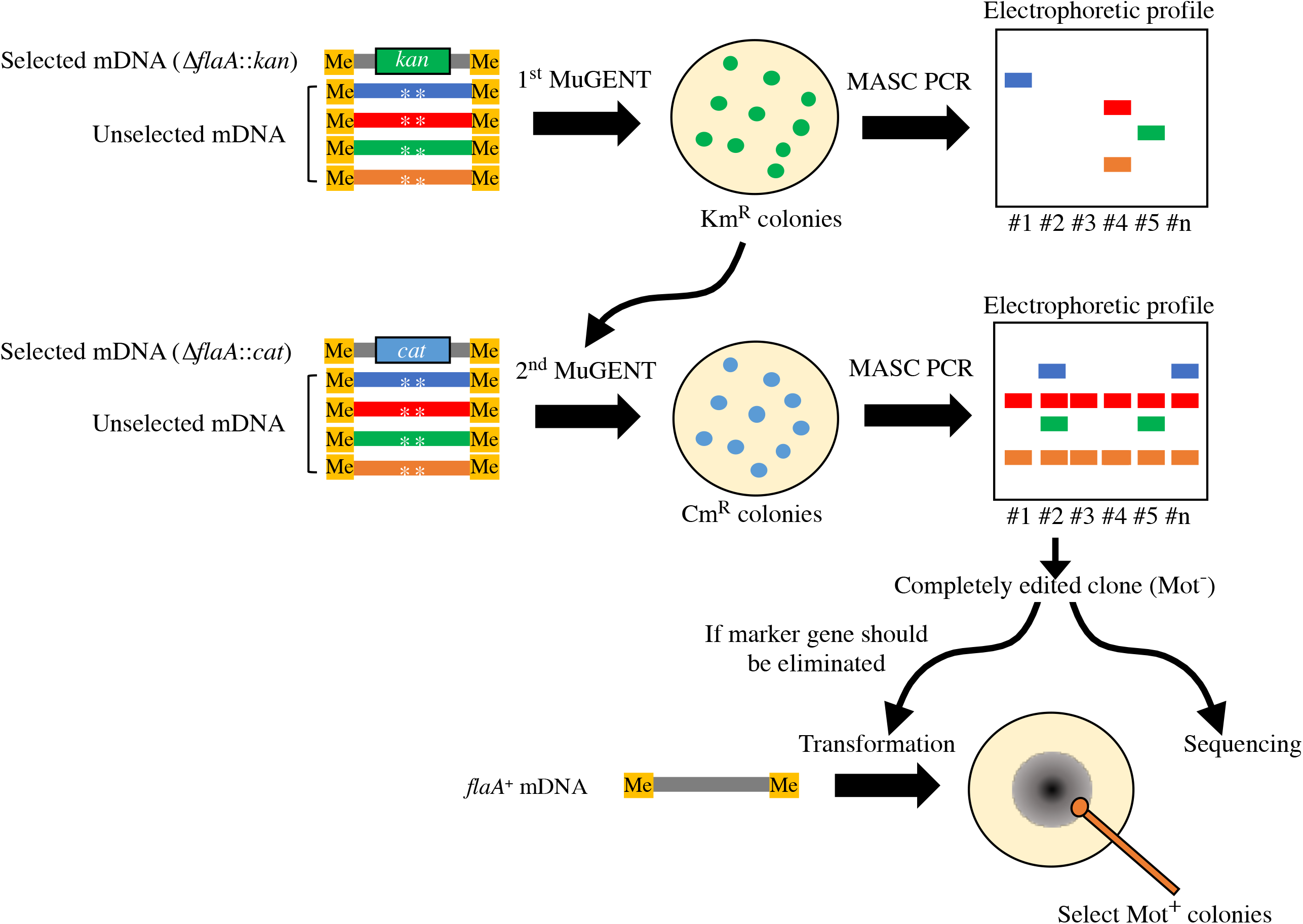
Overview of MuGENT in *C. jejuni.* Multiple unselected mDNA fragments (used for introducing genome edits of interest) were mixed at equimolar concentrations with a selected mDNA fragment, in order to cotransform *C. jejuni* cells. During MuGENT, we swapped different selective markers at the *flaA* gene during every cycle, resulting in the cells becoming nonmotile (Mot^-^). This also enabled easy removal of marker genes and reversion to the wild-type allele by transformation with a *flaA*^+^ mDNA and subsequent selection of motile (Mot^+^) clones. Genome edits in the transformants were verified by MASC PCR and nucleotide sequencing.

To demonstrate the efficacy of MuGENT in *C. jejuni,* we targeted the biosynthetic genes, CPS and LOS. These cell-surface molecules are known to play key roles in interactions that affect bacteriophage infectivity, chick colonization, invasion of human epithelial cells, and host immune responses [34–40]. CPS is also the primary antigenic determinant of the Penner serotyping scheme, a passive slide hemagglutination [41], although other surface molecules, including LOS, may contribute to serotype specificity [42]. Because the CPS and LOS gene clusters contain multiple phase-variable genes that are interrupted by polyG tracts, *C. jejuni* cells can generate structural variations of CPS and LOS, which aids in evading killing by host immune systems or predation by bacteriophages [12,43–46]. In addition, phase-variable expression of CPS and LOS markedly decreases the phenotypic stability of *C. jejuni* cells and, thus, may limit related research, including basic studies, epidemiological surveillance, and vaccine development. For example, the NCTC11168 strain (serotype B, antigenic factor HS2) did not provide reproducible results for Penner serotyping, which frequently changed between typeable and untypeable during subcultures (S1 Fig). Although this unstable phenotype may be attributable to ON/OFF switching of the CPS expression due to phase variation [41,47,48], the mechanism whereby phase variation regulates CPS gene expression to determine the Penner serotype remains unknown.

There are 29 polyG/C tracts in the NCTC11168 genome, eight of which are located within the CPS and LOS gene clusters [10]. Six of these phase-variable genes *(cj1420, cj1421, cj1422, cj1426, cj1429,* and *cj1437*) reside in the CPS cluster, whereas the other genes (*cj1139* and *cj1144*) reside in the LOS cluster, which theoretically gives rise to ~2^8^ different combinations of ON (1) /OFF (0) expression states or more specifically “phasotypes” [4]. For example, a bacterium that has the following expression states — *cj1139*^ON^ *cj1144*^OFF^ *cj1420*^OFF^ *cj1421*^OFF^ *cj1422*^OFF^ *cj1426*^OFF^ *cj1429*^OFF^ *cj1437*^OFF^ (in that order) — would have a binary phasotype coded as 1-0-0-0-0-0-0-0, which can be converted to decimal 128 format. In this study, we defined one of the 2^8^ phasotypes generated by these eight phase-variable genes as a “Penner phasotype (PPT).” Using MuGENT, we tried to “lock” all of the eight ORFs into ON or OFF states, where their polyG tracts were altered to translate into the largest possible ORF (locked-”ON” states) or a smaller incomplete ORF by frameshifting through −1 deletions from the ON states (locked-”OFF” states). For example, *cj1139* encodes a glucosyltransferase that mediates phase variation of LOS epitopes responsible for autoimmunity [13], with G8 being in an ON state and G7 in an OFF state (Fig 4A). To prevent replicative slippage at the polyG tracts, we interrupted the continuous run of G residues without changing the translated amino acids by replacing the last G residue of every G triplet with a different nucleotide (Fig 4A). As an example of phase-locked mutant construction, we introduced locked-OFF mutations into *cj1139, cj1144, cj1420, cj1421, cj1422, cj1426, cj1429,* and *cj1437* by repeating the MuGENT cycle to generate a “PPT0” strain. During MuGENT, genome editing from 40 transformants per cycle was monitored by MASC PCR. After the second cycle, we found that 100% of the population had at least one edit (Fig 4B and S2 Fig). We accomplished all eight edits within the 4th cycle, which took less than 2 weeks to perform (Fig 4B and S2 Fig). Thus, MuGENT works effectively in *C. jejuni* and is useful for rapidly introducing multiple mutations.

**Fig 4.**
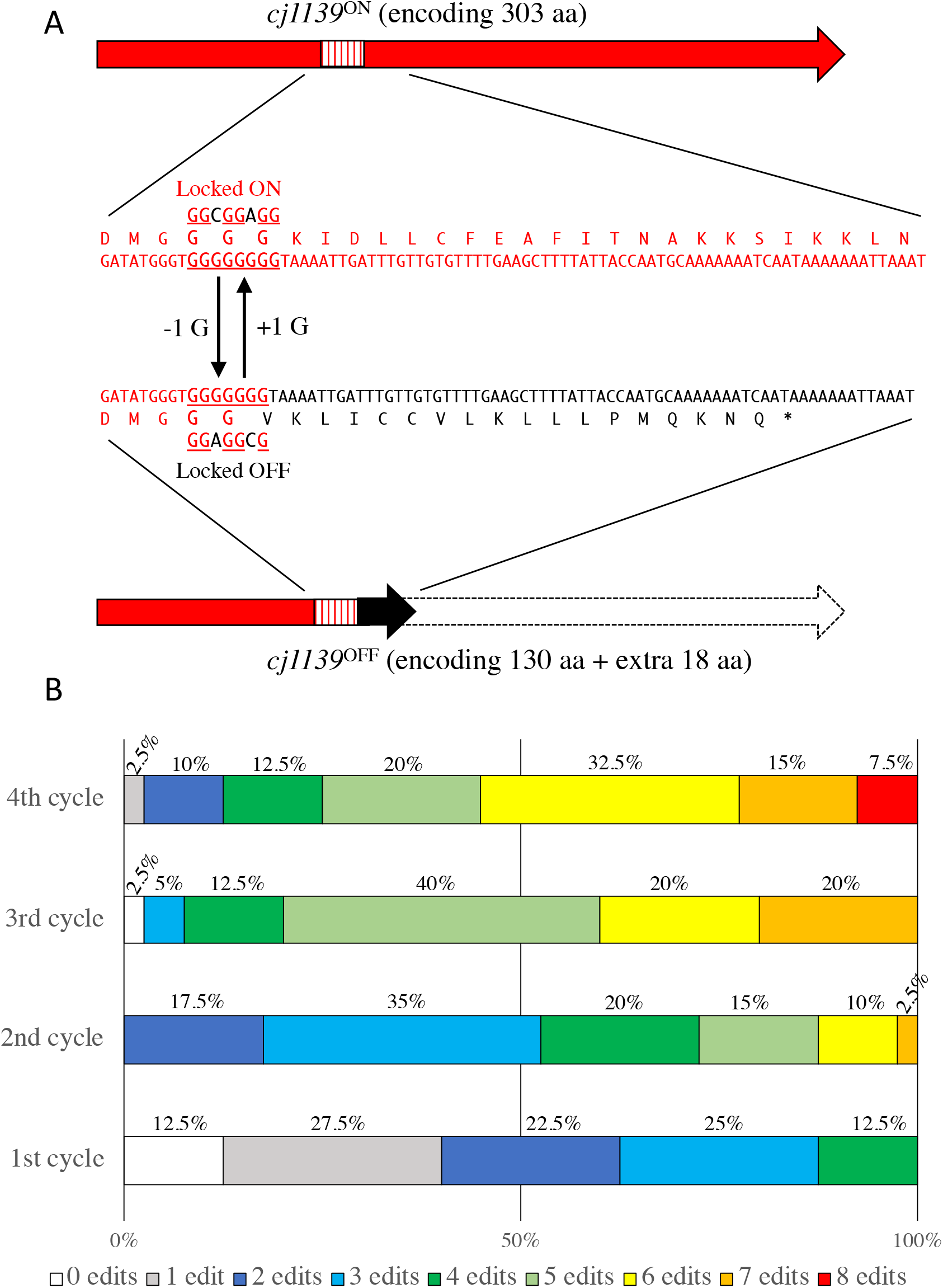
Constructing a strain with all eight phase-variable genes locked into the OFF state by MuGENT against SSR (MuGENT-SSR) sites. The polyG tracts from eight genes (*cj1139, cj1144, cj1420, cj1421, cj1422, cj1426, cj1429,* and *cj1437*) located in the biosynthetic gene clusters for LOS and CPS in NCTC11168 were subjected to MuGENT-SSR to lock them all into OFF states. (A) Example of polyG editing *(cj1139).* The polyG tract was altered to translate into the largest possible ORF (G8: ON state, encoding 303 amino acids [aa]) or a smaller incomplete ORF by introducing frameshifts via −1 deletions (G7: OFF state, encoding 148 amino acids). To prevent replicative slippage at the polyG tract, we interrupted the continuous run of G residues without changing the translated amino acids by replacing the last G residue of every G triplet with a different nucleotide. We also used different nucleotides when locking the genes into ON and OFF states, so that these two different states could be swapped. (B) Distribution of genome edits in the population following successive cycles of MuGENT. This image was based on the MASC PCR data (S2 Fig).

### Improving the phenotypic stability during Penner serotyping using MuGENT

Controlling SSR-mediated phase variation is important for achieving reproducible results with *C. jejuni.* Regarding the low reproducibility of Penner serotyping (S1 Fig), we hypothesized that this may be associated with the phase-variable expression of one or more genes that determine antigenicity. To narrow down candidate genes, we sequenced PPT from naturally occurring typeable and untypeable variants of NCTC11168, termed “PPT-Seq.” PPT-Seq analysis of 12 variants revealed six PPT subtypes, one of which was typeable (Fig 5 and S1 Fig, PPT37). We constructed a strain locked to PPT37 by MuGENT and confirmed that it was also typeable (Fig 5 and S1 Fig). In contrast to the wild-type strain, the locked PPT37 mutant maintained typeability during at least five subcultures (S1 Fig). To evaluate the effect of polyG editing by MuGENT on phase variation, the *astA* gene (encoding arylsulfatase) [49] was fused in frame to the phase-variable gene *cj1426* (Fig 5), which encodes a methyltransferase that methylates the heptose of the CPS repeat unit of NCTC11168 [50], and to its phase-locked derivatives, such that changes in the repeat number would alter arylsulfatase expression [51]. Single colonies grown on BHI agar plates containing 5-bromo-4-chloro-3-indolyl sulfate (XS), a chromogenic substrate of arylsulfatase, were used to measure mutation rates, as described in the Methods section. With the wild-type *cj1426::astA* fusion, the ON-to-OFF mutation occurred at a rate of 2.8 × 10^-2^, whereas the OFF-to-ON mutation rate was 2.3 × 10^-2^ (Table 2). In the locked-ON and locked-OFF constructs, however, switching to different phases was not detectable (Table 2). These results suggest that MuGENT can serve as a feasible and effective approach for stabilizing unstable phenotypes generated by phase variation with a defined set of multiple SSRs.

**Fig 5.**
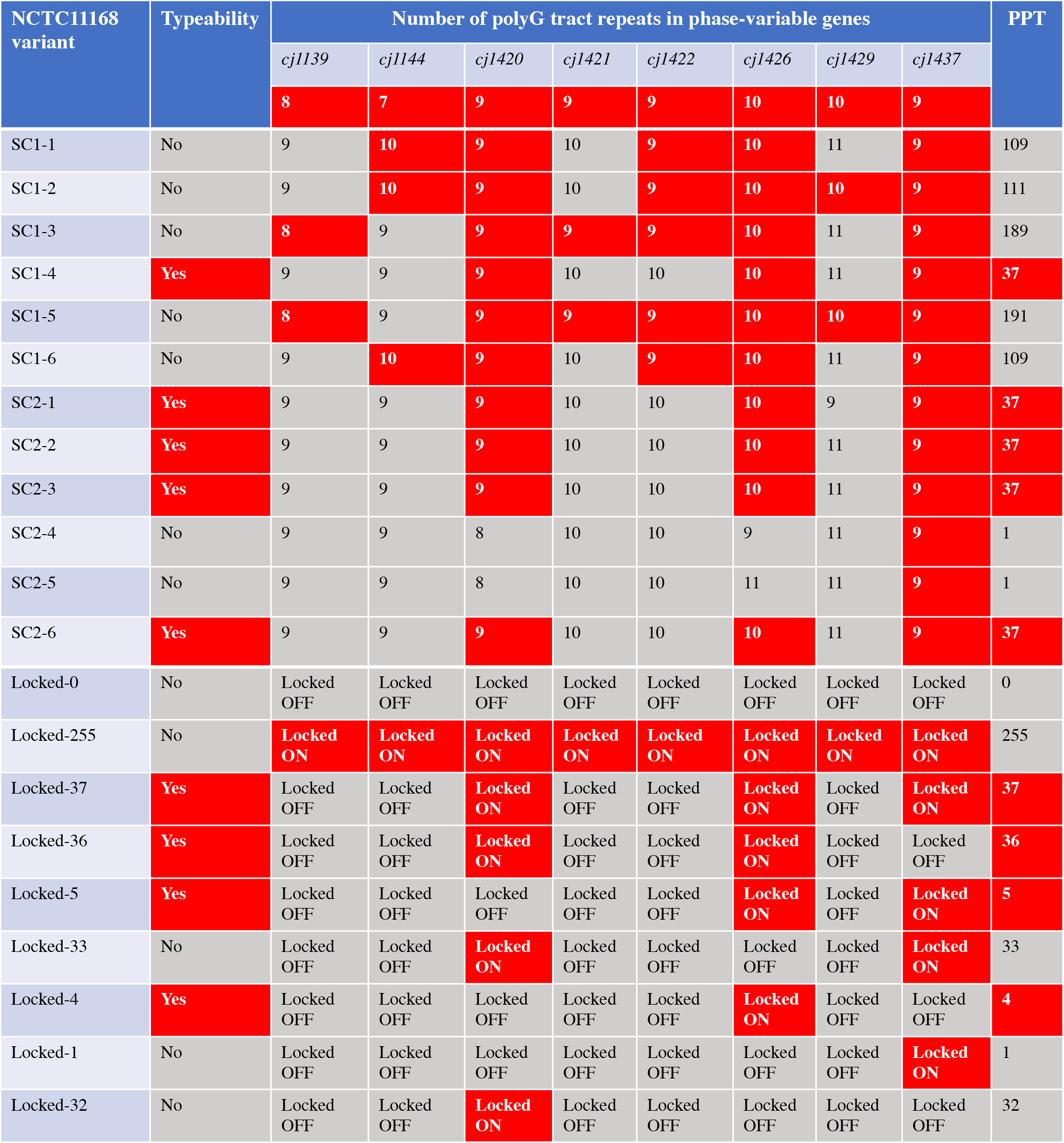
Penner serotyping and phasotyping of naturally occurring and phase-locked variants of NCTC11168. After naturally occurring variants (typeable or untypeable by Penner serotyping; SC1-1 to SC2-6, S1 Fig) were subjected to PPT-Seq, their PPTs were decoded (indicated by decimal). The numbers with in red boxes indicate the numbers of repeat polyG tracts in the ON configurations, whereas those in the gray boxes indicate those in the OFF configurations. The serotyping results for the phase-locked variants are shown in S3 Fig. The following phase-locked strains were used: Locked-0, SYC1P000; Locked-255, SYC1P255; Locked-37, SYC1P037; Locked-36, SYC1P036C; Locked-5, SYC1P005C; Locked-33, SYC1P033C; Locked-4, SYC1P004K; Locked-1, SYC1P001C; Locked-32, SYC1P032K.

**Table 2.**
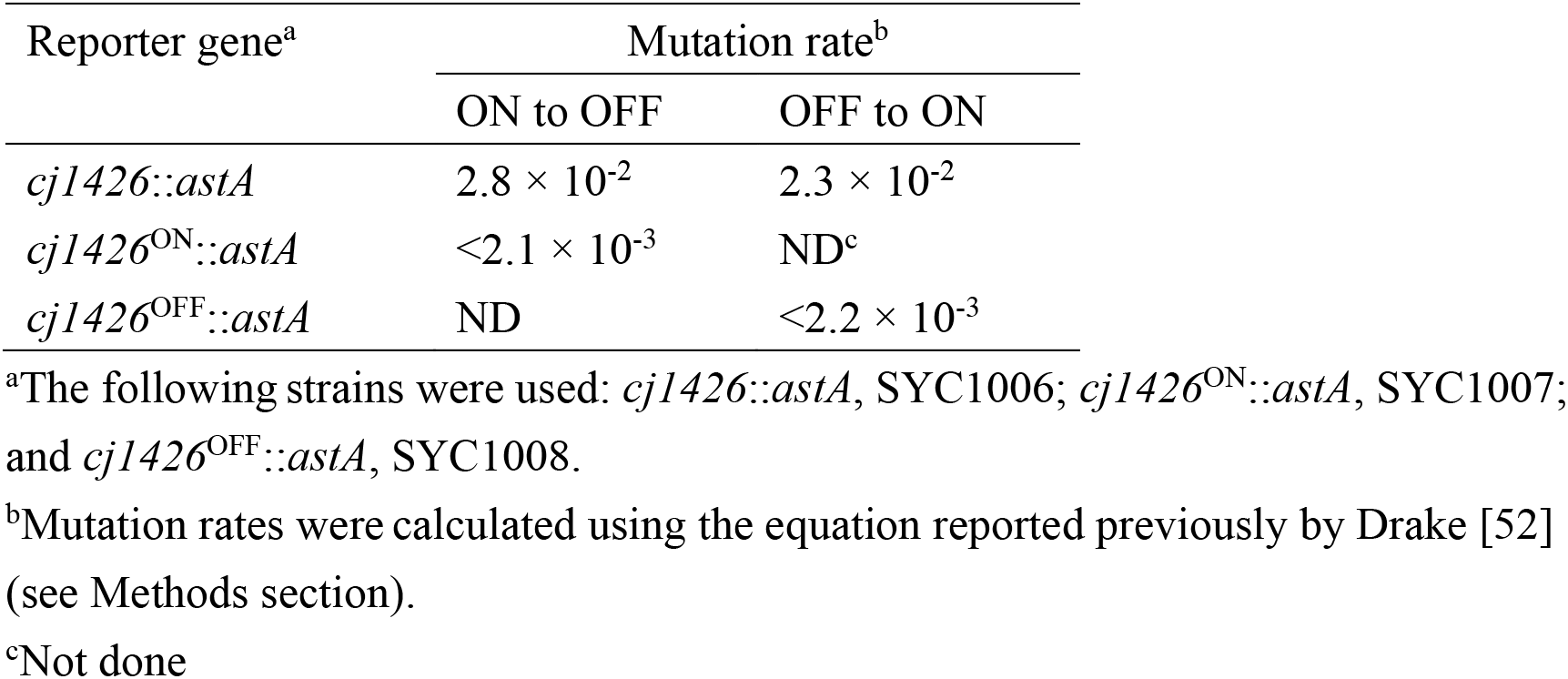
Effect of the polyG editing on phase variation of *cj1426*.

### Using MuGENT-SSR to identify a phase-variable gene that determines the Penner serotype of NCTC11168

Phenotypic engineering using MuGENT-SSR combined with PPT-Seq identified PPT37 as a PPT subtype typeable for Penner serotyping. In PPT37, *cj1420, cj1426,* and *cj1437* were in the ON state, whereas the other five genes were in the OFF state. These findings suggest that one or more genes of these “ON” genes may determine typeability. To elucidate the responsible gene(s), we constructed strains locked to various PPT subtypes by MuGENT-SSR and then performed serotyping. We demonstrated that *cj1426* expression was indispensable and sufficient for serological typing, whereas *cj1420* and *cj1437* expression were not (Fig 5 and S3 Fig). The CPS repeat unit of *C. jejuni* NCTC11168 consists of β-d-ribose, β-D-N-acetylgalactosamine (GalfNAc), α-d-glucuronic acid modified with 2-amino-2-deoxyglycerol at C-6, and 3,6-O-methyl-D-glycero-a-L-gluco-heptose (Hep) [53] (S4 Fig). Examination of the CPS structure by high-resolution magic-angle spinning nuclear magnetic resonance spectroscopy revealed highly variable or no modification of the methyl, ethanolamine, aminoglycerol, and phosphoramidate groups of the repeating unit [38,54]. In particular, changes in methyl (Me) and phosphoramidate modifications were regulated by at least three phase-variable genes located in the CPS cluster (*cj1421*, *cj1422*, and *cj1426*). The *cj1426* gene encodes the methyltransferase involved in 6-O-Me modification of the Hep residue of NCTC11168 CPS [50] (S4 Fig), suggesting that the presence of this modification may be an antigenic determinant of the Penner B serotype (or antigenic factor HS2). However, we observed that some PPT subtypes (including naturally occurring variants PPT109, PPT111, PPT189, and PPT191, and a phase-locked variant PPT255), where *cj1426* and some or all of the other phase-variable genes (including *cj1144, cj1421, cj1422*, and *cj1429*) were simultaneously switched ON, were untypeable (Fig 5, S1 and S3 Figs). Of these genes, the *cj1421* and *cj1422* genes have already been demonstrated to encode O-Me-phosphoramidate (MeOPN) transferases that attach MeOPN to the GalfNAc and Hep residues, respectively [38]. Thus, one or more CPS modifications catalyzed by these gene products may interfere with the binding of specific antibodies to the epitope (S4 Fig). To directly test this possibility, structural analysis using nuclear magnetic resonance spectroscopy and deeper genetic and immunogenic studies are required.

Similar results were reported previously regarding the phase-variable CPS of *C. jejuni.* In a series of studies on the tropism of *Campylobacter* bacteriophages, Sørensen *et al.* demonstrated that the MeOPN-modified GalfNAc of the NCTC11168 CPS acts as a receptor of the F336 phage, but that its receptor function was modulated by the presence or absence of other CPS modifications [37,45]. In addition, a study of *C. jejuni* 81-176 (Penner serotype R, antigenic factors HS23 and HS36) revealed that phase-variable MeOPN modifications at the three CPS galactose residues modulated serum resistance [44]. These previous findings and our current findings suggest that phase-variable changes in the CPS structure of *C. jejuni* occur at two levels: (1) phase variations of the receptors and epitopes themselves, and (2) interference with receptor and epitope functions by chemical modifications at different positions. These combinatorial effects may aid in rapidly avoiding killing by phages and the host immune system, while lowering the typeability of Penner serotyping.

## Concluding remarks

Here, we demonstrate that MuGENT is applicable for genetic engineering of *C. jejuni*. In this genetically unstable species, MuGENT specifically provided a feasible and effective approach for editing multiple hypermutable SSRs. Specifically, MuGENT-SSR was performed to uncover the contributions of multiple phase-variable genes to specific phenotypes. By combining MuGENT-SSR with whole-genome SSR analysis [4,10] may enable comprehensive studies of numerous phase-variable genes (~30 polyG/C-containing genes per genome) in order to decode the “phasotypes” that determine specific phenotypes and more collective behaviors, such as colonization of animal hosts [9,16–20].

We also propose that MuGENT-SSR can be utilized to engineer strains suitable for serological typing and vaccination. In *C. jejuni* NCTC 11168, the majority of polyG/C SSRs are clustered in genomic regions encoding proteins involved in the biosynthesis of cell-surface antigenic determinants, including CPS and LOS [5]. Thus, phase variation makes these antigens less desirable as serodeterminants and vaccine candidates. Penner serotyping is often utilized to investigate epidemiological associations with Guillain– Barré syndrome, but its low typeability is currently problematic [55]. The generation of phase-locked strains by MuGENT-SSR may overcome these defects and stabilize their antigenicity, thereby increasing the supply of stable serotyping antisera and vaccines. Furthermore, PPT-decoding of Penner serotypes provides a reliable approach for identifying antigenic determinant genes and improving or developing DNA-based typing methods that do not rely on CPS expression [56].

## Methods

### Bacterial culture conditions

*C. jejuni* strains used in this study (Table 1) were routinely cultured for 24–48 h at 42°C on BHI (Becton Dickinson) plates containing 1.3% agar (Kyokuto) under microaerophilic conditions using Mitsubishi Anaeropack MicroAero gas generator packs (Mitsubishi Gas). Motility was determined by culturing the cells on BHI plates containing 0.3% agar. For liquid culture, fresh single colonies grown on BHI agar plates were inoculated into 5 ml of BHI broth in 25 ml volumetric test tubes and then cultured overnight at 42°C with reciprocal shaking at 160 rpm in a Taitec Precyto MG-71M-A Obligatory Anaerobe Culture System (Taitec), which can efficiently create a microaerophilic atmosphere by actively aerating gases into each test tube. The aeration conditions in each test tube were set to 5% O_2_, 10% CO_2_, and 85% N2 at a constant flow rate (10 ml/min). The antibiotics used were 25 μg/ml Cm, 50 μg/ml Km, and 10 μg/ml Sm. If necessary, XS (Sigma) was added to the BHI agar plates at a final concentration of 100 μg/ml.

### DNA manipulation

PCR amplification of DNA was performed using a LifeECO thermal cycler (version 1.04, Bioer Technology) and Quick Taq^®^ HS DyeMix DNA polymerase (Toyobo). Sanger sequencing was performed by Fasmac. PCR products were purified using a High Pure PCR Product Purification Kit (Roche). Customized oligonucleotide primers (S1 Table) were purchased from Fasmac and Hokkaido System Science. Chromosomal and plasmid DNA molecules were extracted with a DNeasy Blood and Tissue Kit (Qiagen) and a High Pure Plasmid Isolation Kit (Roche), respectively.

### Construction of *pSYC-cat* and *pSYC-kan*

The pSYC-*cat* and pSYC-*kan* plasmids (Table 1) were constructed by Fasmac as follows: DNA fragments containing the Cm-resistance gene (*cat*) (GenBank accession number M35190.1, nucleotides 1–1,034) and the Km-resistance gene *(kan)* (GenBank accession number M26832.1, nucleotides 1–1,426) from *C. coli* [57,58] were synthesized and cloned into the EcoRV site of pUCFa (Fasmac) to generate pSYC-*cat* and pSYC-*kan*, respectively.

### Natural transformation of *C. jejuni* cells using PCR products

Genetically engineered *C. jejuni* strains were constructed by performing natural transformation [26] optimized to use PCR products as donor DNA. Briefly, the procedures include three processes: (1) PCR amplification of donor DNA with methylation sites, (2) methylation of the amplified DNA, and (3) transformation of *C. jejuni* with methylated DNA.

#### PCR amplification of donor DNA with methylation sites

The donor DNA fragments used for natural transformation were amplified using specific primers and templates. The typical amplified fragment contained an antibiotic-resistance gene (or mutation) with 5’- and 3’-flanking regions homologous to the target site and an EcoRI-recognition sequence (GAATTC) on both sides. DNA was amplified via splicing by overlap extension PCR, which includes a two-step amplification process [59,60]. The first step independently generated two or three fragments with overlapping sequences. In the second step, the splicing by overlap extension PCR fragments amplified in the first step are ligated. The thermal cycling program was as follows: 94°C for 5 min, followed by 30 cycles of 94°C for 30 s, 55°C for 30 s, and 68°C for 1 min per kb. S2 Table shows the specific combinations of template DNA and primer pairs used for PCR amplification.

#### Methylation of donor DNA

DNA methylation was performed using a 25 μl mixture containing 20 nM DNA, EcoRI methyltransferase reaction buffer (New England Biolabs; 50 mM Tris-HCl, 50 mM NaCl, 10 mM ethylenediaminetetraacetic acid, pH 8.0), 80 μM *S*-adenosylmethionine (New England Biolabs), and 40 units of EcoRI methyltransferase (New England Biolabs). After incubation for 2.5 h at 37°C, the reaction was terminated by incubation for 20 min at 65°C. Methylated DNA was purified using the High Pure PCR Product Purification Kit and dissolved in 100 μl H_2_O.

#### Natural transformation

Overnight cultures of *C. jejuni* cells were diluted 1/50 in 5 ml BHI broth and grown until they reached an optical density and 600 nm (OD_600_) of approximately 0.15. Each culture (900 μl) was mixed with methylated DNA (100 μl) and then statically incubated overnight in a 25 ml volumetric plastic test tube. On the following day, 10-fold serial dilutions of the culture were spread onto BHI agar plates, with or without antibiotics. The transformation frequency was defined as the number of antibiotic-resistant CFUs divided by the total number of CFUs.

### Evaluation of natural cotransformation in *C. jejuni*

Natural cotransformation in *C. jejuni* was examined using two different PCR fragments, Δ*flaA::kan* and *rpsL*^+^ (see S2 Table). The former fragment (Δ*flaA::kan-1* or Δ*flaA::kan*-2) had 1,000-bp regions of homology, whereas the latter had 1,000-bp or 2,000-bp regions of homology (*rpsL*^+^-8-1, *rpsL*^+^-9-1, *rpsL*^+^-8-2, or *rpsL*^+^-9-2). The Δ*flaA::kan* and *rpsL*^+^ fragments were independently methylated and then mixed for purification as described above. After transformation of SY1003 and SY2003 with these methylated DNA fragments, the reactions were spread onto BHI agar plates containing Km. One hundred Km^R^ transformants were replica plated onto BHI agar plates containing Sm. The cotransformation frequency (%) was calculated as follows: 100 × number of Sm^S^ CFUs in 100 Km^R^ CFUs.

### MuGENT-SSR

MuGENT-SSR was performed with *C. jejuni* cells, as follows. After independently methylating multiple unselected fragments, including eight fragments used for editing of the eight polyG sites in the biosynthetic LOS and CPS gene clusters in NCTC11168, as well as a single selected fragment (either Δ*flaA::kan-1* or Δ*flaA::cat*-1), a mixture of these fragments was prepared as described above. Unselected fragments had 1,500–2,000-bp regions of homology, whereas the selected fragment had a 1,000-bp region of homology (S2 and S3 Tables). After transformation with the DNA mixture, the bacteria were plated onto BHI agar plates containing the antibiotic corresponding to the resistance protein encoded in the selected fragment. Colonies on selective agar plates were used for MASC PCR (described below). The remaining colonies (3,000–5,000 CFUs) were suspended in BHI broth and diluted in 5 ml BHI broth to an OD_600_ of 0.05. After the cells were grown to an OD_600_ of 0.15, the next cycle of MuGENT was performed using unselected fragments and a selected fragment with a different resistance gene from that used in the previous cycle. MuGENT cycles were repeated until editing was completed. Genome editing was ultimately verified by nucleotide sequencing using specific primers (see the PPT-Seq section below). If necessary, the Δ*flaA* mutation maintained during MuGENT was reverted to the wild-type allele by transformation using the *flaA*^+^ fragment with a 1,000-bp region of homology. After the transformation, colonies showing motility (and sensitivity to the antibiotic used for selection) were chosen for further study.

### MASC PCR

During each cycle of MuGENT, 40 colonies were inoculated in 100 μl of BHI broth in a 96-wel plate and incubated overnight. Each PCR mixture (20 μl) contained 2 μl of the bacterial culture, 2 μL of 2.5 μM primer mix (one of four primer mixes, including Mix ON1, Mix ON2, Mix OFF1, and Mix OFF2; see S4 Table), 6 μl of H2O, and 10 μl of Quick Taq^®^ HS DyeMix. The following thermal cycling program was used for MASC PCR: 94°C for 5 min, followed by 35 cycles of 94°C for 30 s, 62.6°C for 30 s, and 68°C for 3 min. If necessary, several single colonies were isolated from positive colonies (to eliminate contamination by unedited clones) and further assessed for genome editing by MASC PCR.

### PPT-Seq

The eight phase-variable genes located in the LOS- and CPS-biosynthetic gene clusters in NCTC11168 were amplified by PCR, and the repeat numbers of the polyG SSR tracts were determined by nucleotide sequencing. S5 Table shows the specific combinations of target genes and primers used for PPT-Seq.

### Penner serotyping

A single colony was streaked on the surface of a horse blood agar plate (Kyokuto) and incubated for 48 h at 42°C to determine the Penner serotype [61]. After the resulting colonies grown on the plate were suspended in 250 μL of 0.9% NaCl, serotyping was performed using *Campylobacter* antisera (Denka), including a commercial 25 antisera set, and a reagent for preparing sensitized blood cells (Denka) according to the manufacturer’s instructions. Phenotypic stability was assessed by repeating the subculture cycle and serotyping. Serotyping was performed using six single colonies for each cycle.

### Measurement of mutation rates

The mutation rates of the phase-variable *cj1426* gene and its phase-locked variants were determined as described previously [62], with some modifications. *C. jejuni* strains carrying *astA* to *cj1426* translational fusions were streaked on BHI agar plates containing XS. During growth on the medium, blue colonies were in the *cj1426*-ON phase, whereas white colonies were in the *cj1426*-OFF phase. Single blue or white colonies were picked using a micropipette tip were resuspended in 500 μL BHI, after which 250 μl of 10^4^-, 10^5^-, and 10^6^-fold dilutions were spread onto BHI agar plates containing XS. The number of variant colonies that switched to different phases and the total numbers of colonies were counted. Ten independent single colonies were examined for each strain and phase. To estimate the mutation rate, the total number of colonies was averaged for all single colonies tested, and the median value for the frequency of variants per colony was calculated. The mutation rate was calculated using the following equation: *μ* = 0.4343*f*/*log*(*Nμ*), where *μ* is the mutation rate, *f* is the median frequency, and *N* is the average population size [52]. The *μ* value was determined by solving the equation using the Goal Seek function in Microsoft Excel.

## Acknowledgments

We thank Jiro Mitobe for helpful suggestions.

## Supporting information captions

**S1 Fig. Penner serotyping of naturally occurring and phase-locked NCTC11168 variants.** Phenotypic stability was examined during successive subcultures, which highlights only the results obtained using anti-serotype B antiserum. Red-numbered colonies were typeable, and black-numbered colonies were untypeable.

**S2 Fig. MASC PCR of transformants following successive cycles of MuGENT-SSR.**

**S3 Fig. Penner serotyping of phase-locked NCTC11168 variants.**

**S4 Fig. Putative modification patterns of the CPS repeat unit in naturally occurring and phase-locked NCTC11168 variants.** Figures were modified from illustrations published in Sternberg et al, J Mol Biol 2013 (425) 186-197.

**S1 Table. Primers used in this study.**

**S2 Table. Specific combinations of template DNA and primers used to amplify donor DNA templates.**

**S3 Table. Specific combinations of donor DNA molecules and recipient strains used for natural transformation.**

**S4 Table. Primer mixes used for MASC PCR.**

**S5 Table. Specific combinations of target genes and primers in PPT-Seq.**

**S6 Table. Primer sets used for allele-specific PCR and sequencing of *cj1426::astA* translational fusions.**

