## Supplementary material for "Stabilizing genetically unstable simple sequence repeats in the *Campylobacter jejuni* genome by multiplex genome editing: a reliable approach for delineating multiple phase-variable genes": S1 Fig

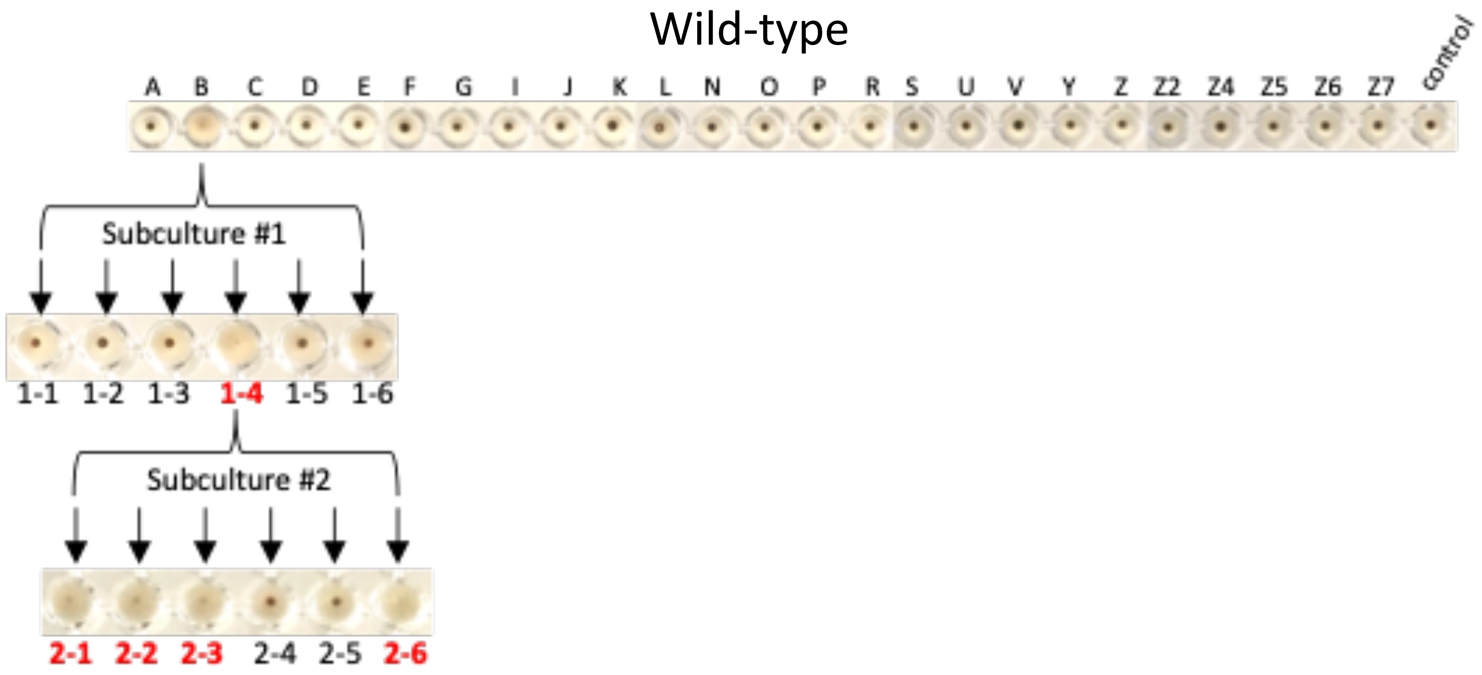


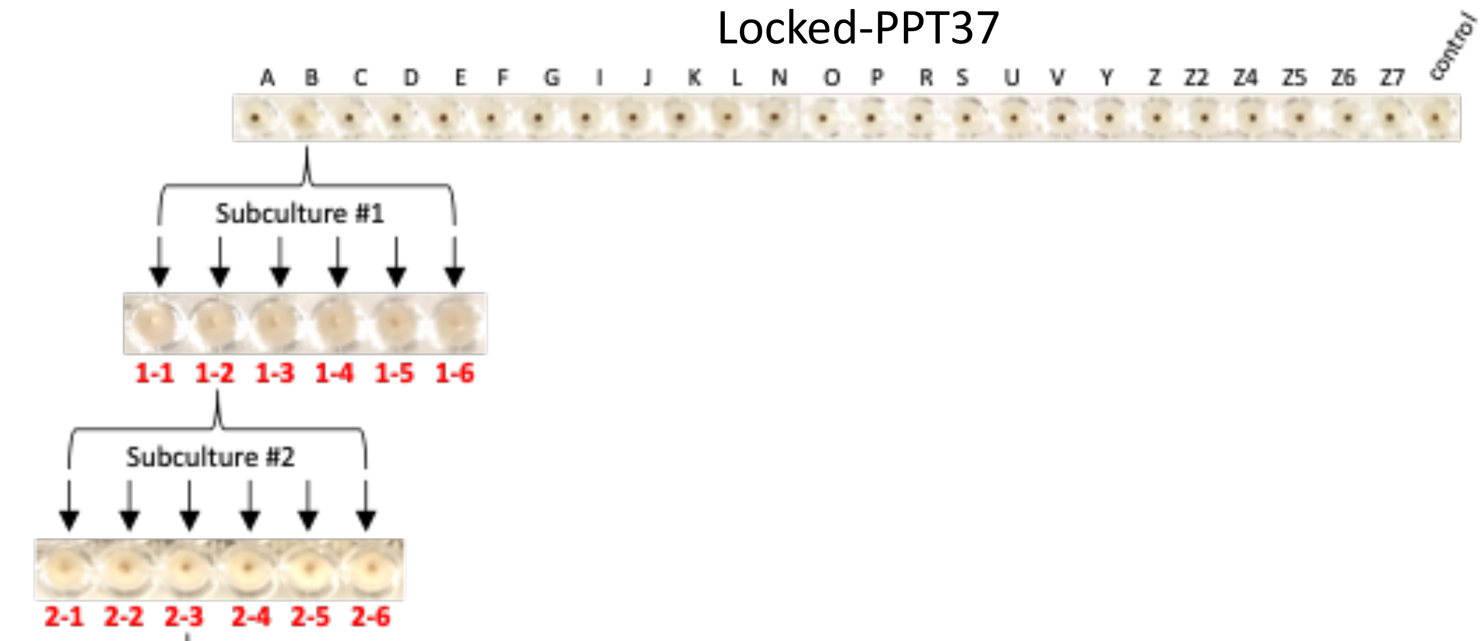


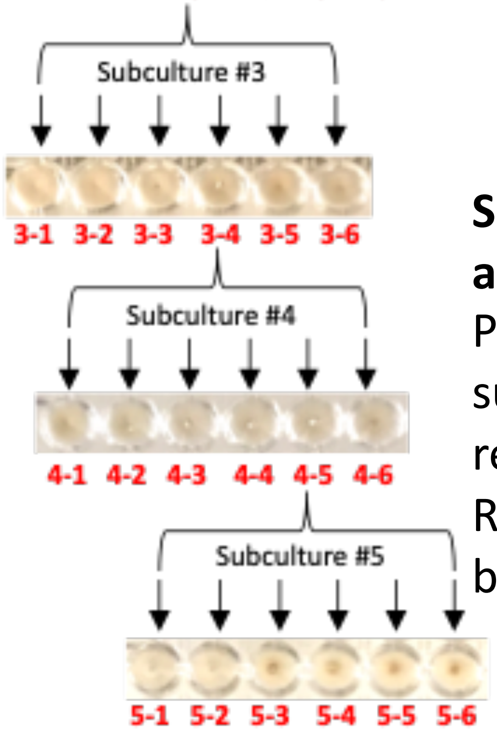


**S1 Fig. Penner serotyping of naturally occurring and phase-locked NCTC11168 variants.** Phenotypic stability was examined during successive subcultures, which highlights only the results obtained using anti-serotype B antiserum. Red-numbered colonies were typeable, and black-numbered colonies were untypeable.
