## Supplementary figures and images for "Stabilizing genetically unstable simple sequence repeats in the *Campylobacter jejuni* genome by multiplex genome editing: a reliable approach for delineating multiple phase-variable genes"

### S2 Fig

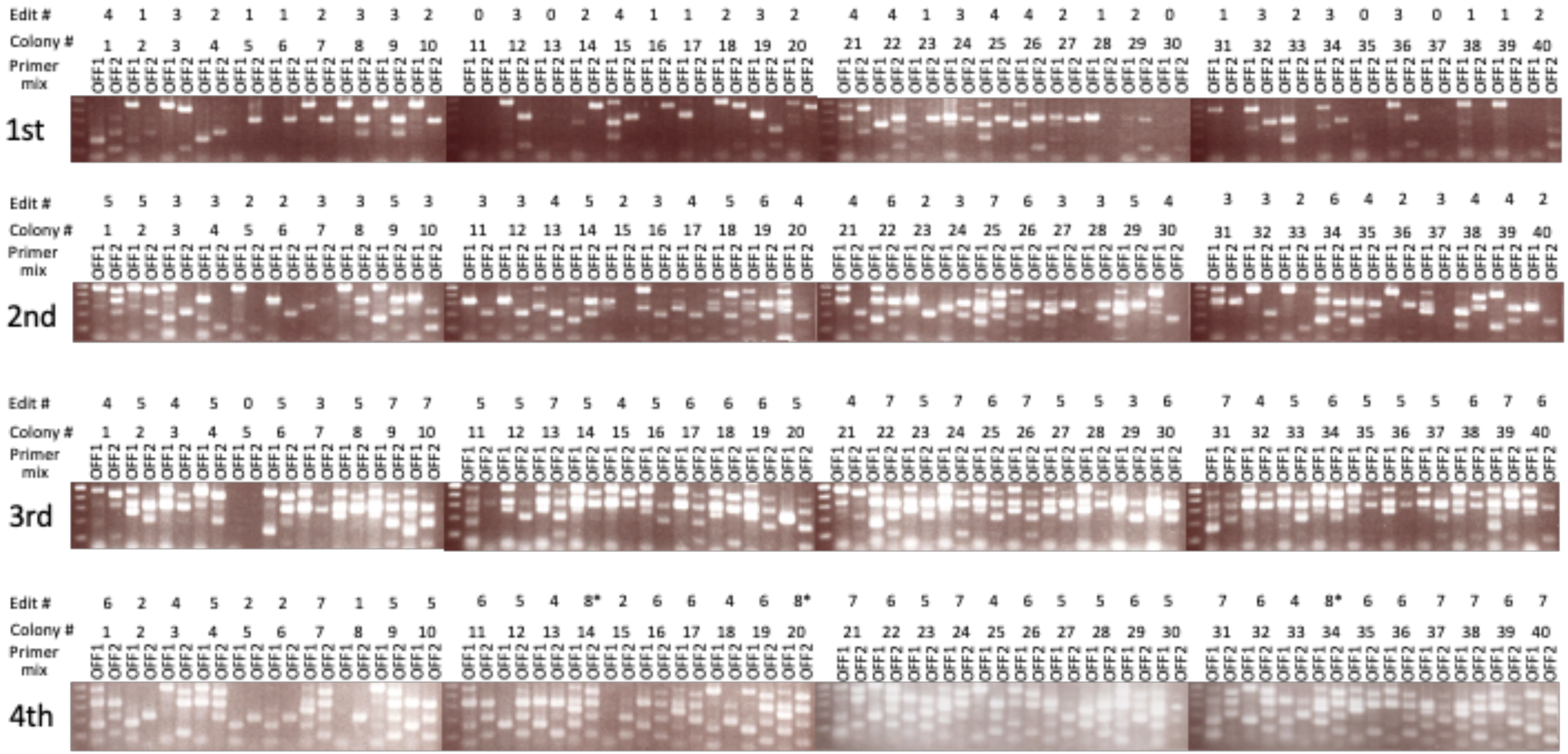


**S2 Fig. MASC PCR of transformants following successive cycles of MuGENT-SSR.**

### S3 Fig

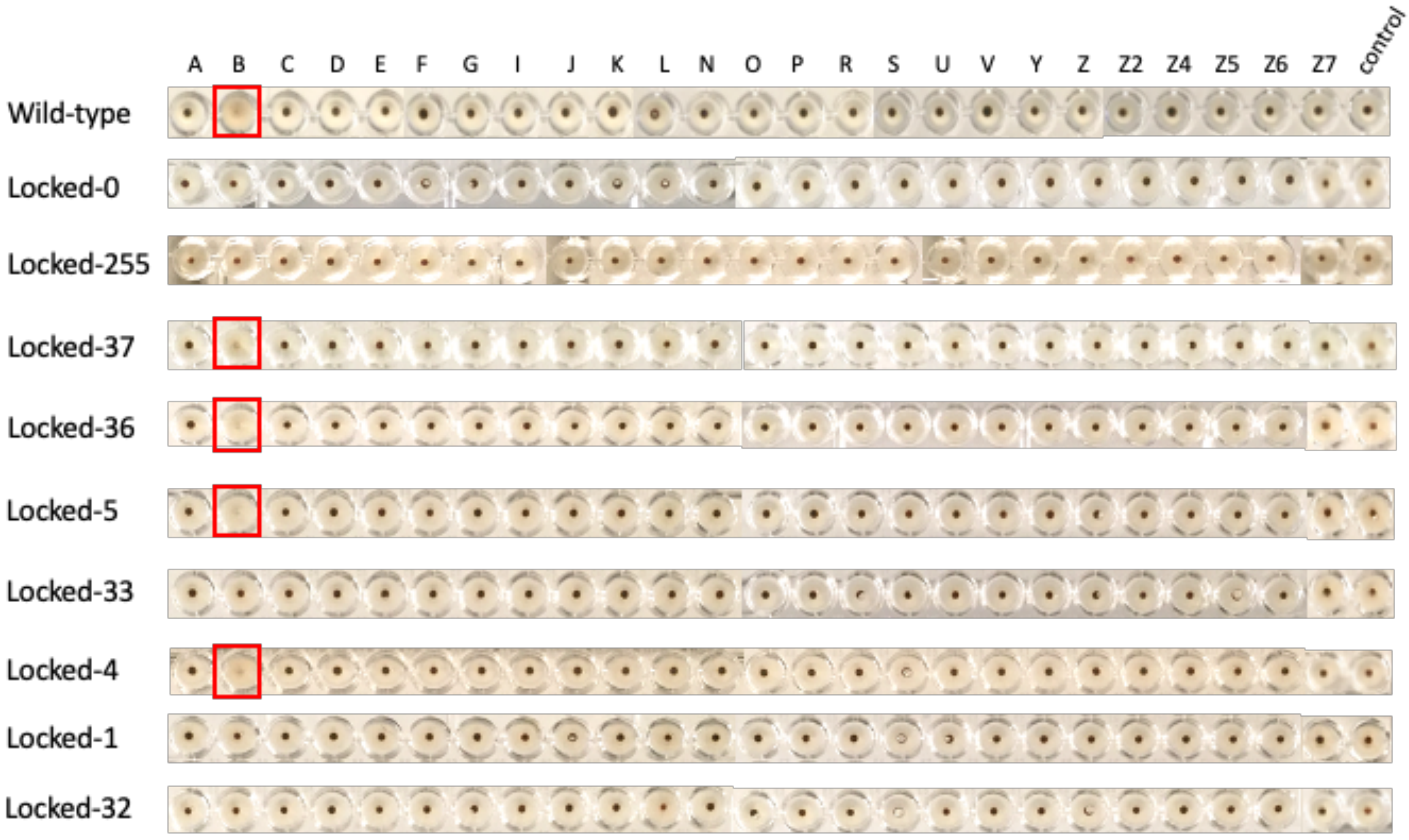


**S3 Fig. Penner serotyping of phase-locked NCTC11168 variants**
