## Supplementary material for "Stabilizing genetically unstable simple sequence repeats in the *Campylobacter jejuni* genome by multiplex genome editing: a reliable approach for delineating multiple phase-variable genes": S4 Fig

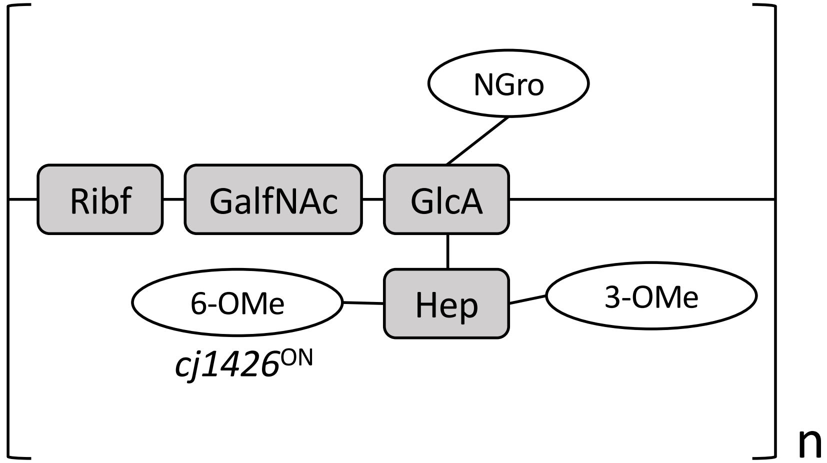


PPT4, PPT5, PPT36, and PPT37
Typeability: yes

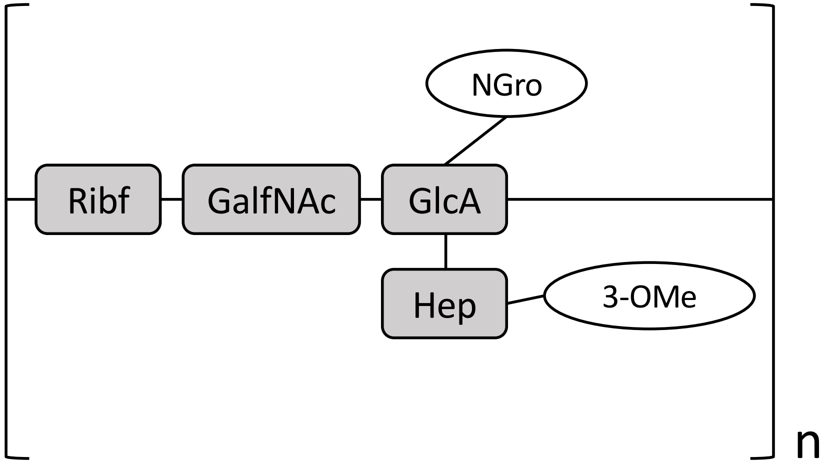


PPT0, PPT1, PPT32, and PPT33
Typeability: no

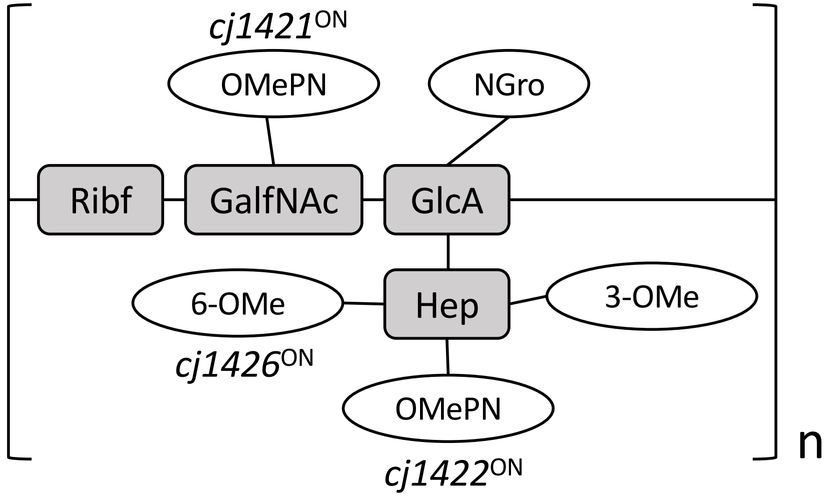


PPT189, PPT191, and PPT255
Typeability: no

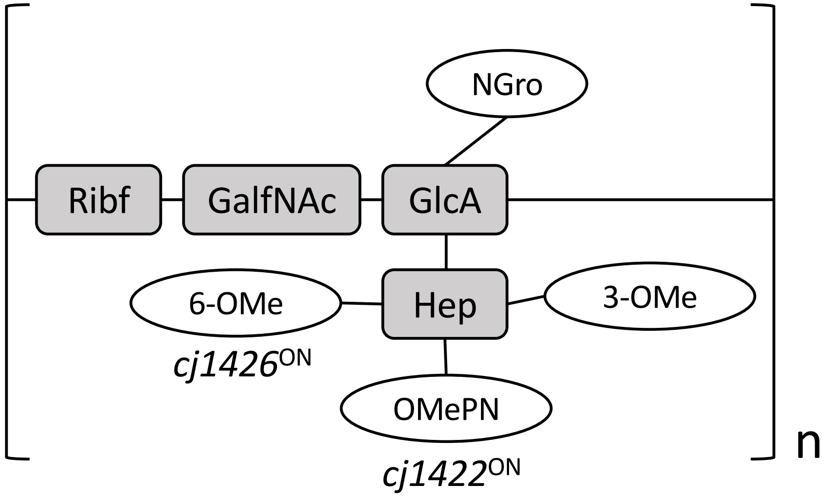


PPT109 and PPT111
Typeability: no

**S4 Fig. Putative modification patterns of the CPS repeat unit in naturally occurring and phase-locked NCTC11168 variants.** Figures were modified from illustrations published in Sternberg *et al*, *J Mol Biol* 2013 (425) 186-197.
