## Supplementary material for "Stabilizing genetically unstable simple sequence repeats in the *Campylobacter jejuni* genome by multiplex genome editing: a reliable approach for delineating multiple phase-variable genes": S1 Table

**S1 Table. Primers used in this study**

| Name | Sequence[a] (5′–3′) |
|---|---|
| Used for natural transformation | |
| 176_1669-f1000E | GGGGAATTCCAACTAAGGTACTTTCGTCC |
| 176_1669-pUCFa-r1 | GGAAGAGCGTCGTTAAGGGAAAGGCAGCGTCTAGAGATTTTC |
| 176_1669-r1000E | GGGGAATTCGGTTTAATAAGCACAGCATTTGC |
| 176rpsLmt-f1000 | GCGTTCTAAAGAAAGAATTATTATCCAAGC |
| 176rpsLmt-f1000E | GGGGAATTCGCGTTCTAAAGAAAGAATTATTATCCAAGC |
| 176rpsLmt-f100E | GGGGAATTCGTAAGACTTACTAGTGGCTTTG |
| 176rpsLmt-f2000E | GGGGAATTCCGGCGATTTTACTTGTGGAG |
| 176rpsLmt-f500 | TAGCGGGTAAATTTGACTACTTAGAAG |
| 176rpsLmt-f500E | GGGGAATTCTAGCGGGTAAATTTGACTACTTAGAAG |
| 176rpsLmt-f50E | GGGGAATTCTCATAACTTGCAAGAACACAGC |
| 176rpsLmt-r1000 | TGAACAAAATACCGCAACAGCACCATC |
| 176rpsLmt-r1000E | GGGGAATTCTGAACAAAATACCGCAACAGCACCATC |
| 176rpsLmt-r100E | GGGGAATTCGCGTTTAGCACCATATTTAGAACG |
| 176rpsLmt-r2000E | GGGGAATTCCATCTGTAGAAACTCTAAAACTTGG |
| 176rpsLmt-r500 | AACGAATCGCCAAAGCTTGTTGTCTAGC |
| 176rpsLmt-r500E | GGGGAATTCAACGAATCGCCAAAGCTTGTTGTCTAGC |
| 176rpsLmt-r50E | GGGGAATTCCTGTATCAAGAGCACCACGAAC |
| 68rpsLmt-f1000 | GCGATTGTAGAATGTGGTGGAAAAATCAC |
| 68rpsLmt-f1000E | GGGGAATTCGCGATTGTAGAATGTGGTGGAAAAATCAC |
| 68rpsLmt-f2000E | GGGGAATTCGAAAGTTTATCCCACATAATGGAG |

| | |
|---|---|
| 68rpsLmt-f500 | AGTAATTTCTTAAAATCAATTTTGG |
| 68rpsLmt-f500E | GGGGAATTCAGTAATTTCTTAAAATCAATTTTGG |
| astA-cj1426c-f2 | GCAATGCCTTTTAGTGTTGATCAAGCATTCAATCCTAAAAAATAATCAGAGAAAA AGATAGGGGAG |
| astA-f2 | AGACTTAGCAAAACTCTTTGTATGGCACTTTTGGCGGGC |
| astA-r2 | TTATTTTTTAGGATTGAATGCTTGATCAACACTAAAAGGCATTGC |
| c-cat-f1 | ATTCCCACAACGCCGGAAAC |
| c-cat-r2 | AATGAAGCTCCGCAGGACGC |
| c-kan-f1 | GATAAACCCAGCGAACCATT |
| c-kan-r1 | GCTTTTTAGACATCTAAATCTAGG |
| cat-cj1339c(flaA)-f2 | GCGTCCTGCGGAGCTTCATTGCTGCAATATATACAAATCC |
| cat-flaA81176-f2 | GCGTCCTGCGGAGCTTCATTGCAATGGCTCAAGCAAATTC |
| cj1139c-OFF(-1)-f2 | GGATATGGGTGGAGGCGTAAAATTGATTTGTTG |
| cj1139c-OFF(-1)-r2 | CAACAAATCAATTTTACGCCTCCACCCATATCC |
| cj1139c-ON-f1 | GGATATGGGTGGCGGAGGTAAAATTGATTTGTTG |
| cj1139c-ON-r1 | CAACAAATCAATTTTACCTCCGCCACCCATATCC |
| cj1139c-f2E | GGGGAATTCTAATCTTACTAAGTGCGATGG |
| cj1139c-r2E | GGGGAATTCAGGTGAAGCTTGATTTTTACC |
| cj1145c-OFF(-1)-f2 | CTTTATCTTAAAAAAAAAGGAGGCTATGGGTAGATCTTG |
| cj1145c-OFF(-1)-r2 | CAAGATCTACCCATAGCCTCCTTTTTTTTTAAGATAAAG |
| cj1145c-ON-f1 | CTTTATCTTAAAAAAAAAGGCGGAGTATGGGTAGATCTTG |
| cj1145c-ON-r1 | CAAGATCTACCCATACTCCGCCTTTTTTTTTAAGATAAAG |
| cj1145c-r1E | GGGGGAATTCTCTCCTTTAAATCCATGGGC |

| | |
|---|---|
| cj1145c-f1E | GGGGGAATTCAGGTGCATAGGTCCACTGTC |
| cj1339c(flaA)-f1E | GGGGGAATTCTGCTACGCATCCTAATATCG |
| cj1339c(flaA)-r2E | GGGGAATTCCATAGGCTACTTGACCTATAG |
| cj1339c(flaA)-cat-r1 | GTTTCCGGCGTTGTGGGAATCAAGCTCATCCATGAACTTG |
| cj1339c(flaA)-kan-r1 | AATGGTTCGCTGGGTTTATCCAAGCTCATCCATGAACTTG |
| cj1420c-OFF(-1)-f2 | CGTATATTGACAGGAGGCGGTATTTTACTGCGATTTGGA |
| cj1420c-OFF(-1)-r2 | TCCAAATCGCAGTAAAATACCGCCTCCTGTCAATATACG |
| cj1420c-ON-f1 | CGTATATTGACAGGTGGAGGCTATTTTACTGCGATTTGGA |
| cj1420c-ON-r1 | TCCAAATCGCAGTAAAATAGCCTCCACCTGTCAATATACG |
| cj1420c-f1E | GGGGAATTCCATTTATCTTACATGATAGG |
| cj1420c-r1E | GGGGAATTCATACGCCCAGATATTATCCG |
| cj1421/22c-OFF(-1)-f2 | GAACATAGACATAACGGAGGCGGTATATAGCATTA |
| cj1421/22c-OFF(-1)-r2 | TAATGCTATATACCGCCTCCGTTATGTCTATGTTC |
| cj1421/22c-ON-f1 | GAACATAGACATAACGGTGGAGGCTATATAGCATTA |
| cj1421/22c-ON-r1 | TAATGCTATATAGCCTCCACCGTTATGTCTATGTTC |
| cj1421c-r1E | GGGGGAATTCCTAAATATCACCATCCAACTCCTTGC |
| cj1422c-f1E | GGGGAATTCATGTACCAAGTGGTAGTGGCTTGGG |
| cj1422c-f2E | GGGGAATTCGTTGGTGTTGTGCTTATTGGTG |
| cj1422c-r2E | GGGGAATTCCCAATACATCGTCTACTTTCACTTC |
| cj1426c-astA-r2 | GCCCGCCAAAAGTGCCATACAAAGAGTTTTGCTAAGTCTATCAGTAATTAAGCCT GCATGTCCAC |
| cj1426c-OFF(-1)-f2 | GTCGATAAATATGGAGGCGGGATGGATATCGTCC |
| cj1426c-OFF(-1)-r2 | GGACGATATCCATCACCGCCTCCATATTTATCGAC |

| | |
|---|---|
| cj1426c-ON-f1 | GTCGATAAATATGGTGGAGGCGATGGATATCGTCC |
| cj1426c-ON-r1 | GGACGATATCCATCGCCTCCACCATATTTATCGAC |
| cj1426c-WT-f1 | GTCGATAAATATGGGGGGGGGATGGATATCGTCC |
| cj1426c-WT-r1 | GGACGATATCCATCCCCCCCCCATATTTATCGAC |
| cj1426c-f1E | GGGGAATTCTGGAAAATCAAGAGTCTTACCC |
| cj1426c-r1E | GGGGAATTCCATTCATAGCCCCGCCACTAGC |
| cj1429c-OFF(-1)-f2 | GATGTATAATGGAGGCGGGATATGAGTGATATTAATGC |
| cj1429c-OFF(-1)-r2 | GCATTAATATCACTCATATCCCGCCTCCATTATACATC |
| cj1429c-ON-f1 | GATGTATAATGGTGGAGGCGATATGAGTGATATTAATGC |
| cj1429c-ON-r2 | CGCCTCCACCATTATACATCACATAACCAC |
| cj1429c-f3E | GGGGAATTCGGTGGTATTCAACTCAAGGTGATCTTAGTG |
| cj1429c-r1E | GGGGAATTCATCCCATATTCTGAGATAGGACGCAAG |
| cj1437c-OFF(-1)-f3 | CCTTAACTACTGGCGGAGGCGGATATTCAATGATTCTGT |
| cj1437c-OFF(-1)-r3 | ACAGAATCATTGAATATACCGCCTCCGCCAGTAGTTAAGG |
| cj1437c-ON-f1 | CCTTAACTACTGGCGGTGGAGGGATATTCAATGATTCTGT |
| cj1437c-ON-r1 | ACAGAATCATTGAATATCCCTCCACCGCCAGTAGTTAAGG |
| cj1437c-r1E | GGGGAATTCGGTACAACAATAGAAACTTTTGGAG |
| cj1437c-f1E | GGGGAATTCGAAAGGATTAAATGGCAATTATC |
| cj1673c-f1000E | GGGGAATTCGCAACTAAGGTACTTTCATC |
| cj1673c-r1000E | GGGGAATTCGGTTTGATAAGTACAGCATTTG |
| cjj81176_1339-f1E | GGGGAATTCTCGGCAAGTACTCATCCTAG |
| cjj81176_1339-r1E | GGGGAATTCTTGGCCGTTATTATCACCATC |
| flaA81176-cat-r1 | GTTTCCGGCGTTGTGGGAATCGAAATCCCATTTTAAATCC |

| | |
|---|---|
| flaA81176-kan-r1 | AATGGTTCGCTGGGTTTATCCGAAATCCCATTTTAAATCC |
| kan-cj1339c(flaA)-f1 | CCTAGATTTAGATGTCTAAAAAGCGCTGCAATATATACAAATCC |
| kan-flaA81176-f1 | CCTAGATTTAGATGTCTAAAAAGCGCAATGGCTCAAGCAAATTC |
| pUCFa-176_1669-f1 | GGAAGAGCACACGTCTGAACTCGCGAAGATGACGAAGGAGAAG |
| pUCFa-f1 | TTTCCCTTAACGACGCTCTTCC |
| pUCFa-r1 | GAGTTCAGACGTGTGCTCTTCC |
| rpsL(CJ0491)-StmR-F2 | GGTGGTAGGGTAAGAGACTTACCAGGGG |
| rpsL(CJ0491)-StmR-R2 | CCCCTGGTAAGTCTCTTACCCTACCACC |

Used for MASC PCR

| | |
|---|---|
| astA-MASCR1 | GCGGTCAAAGGAGACAATCCATAAGG |
| cj1139c-MASCmF2M | TATTAAAATTTTGGATATGGGTGGCGcA |
| cj1139c-MASCmF3M | TATTAAAATTTTGGATATGGGTGGAGcC |
| cj1139c-MASCR2 | TATCTTTTTTGATTATTTTAGCCCACATTGTCC |
| cj1145c-MASCmF1M | TTTTAGATACAATTTACTTTATCTTAAAAAAAAAGGCGcA |
| cj1145c-MASCmF2M | TTTTAGATACAATTTACTTTATCTTAAAAAAAAAGGAGcC |
| cj1145c-MASCR1 | GGATGTTGTGATTCTTGATTTTTTATTATCTTCATCCAC |
| cj1420c-MASCmF1M | GAATTTAATCGTATATTGACAGGTGGAGcC |
| cj1420c-MASCmF2M | GAATTTAATCGTATATTGACAGGAGcC |
| cj1420c-MASCR1 | CTCCTTTTCAATTTCATCAAAAACCGAACC |
| cj1421c-MASCF1 | TGAGGAATTGGTTTACATCAAGCAAC |
| cj1421c/22c-MASCmR1M | GAGTTTTTTAAATAATGCTATATAGCCTCgA |
| cj1421c/22c-MASCmR2M | GAGTTTTTTAAATAATGCTATATACCGCgT |

TATTAAAAT

| | |
|---|---|
| cj1422c-MASCF4 | ATGATTTTGATAGATACGGCACAGTAAATG |
| cj1426c-MASCmF1M | ATGCTTTATGTCGATAAATATGGTGGAGcC |
| cj1426c-MASCmF2M | ATGCTTTATGTCGATAAATATGGAGcC |
| cj1426c-MASCwF1M | CTTTATGTCGATAAATATGGGGGGGcG |
| cj1426c-MASCR1 | TCCGTCTGACTGTCTTGTACACTTTC |
| cj1429c-MASCmF1M | GTGGTTATGTGATGTATAATGGTGGAGcC |
| cj1429c-MASCmF2M | GTGGTTATGTGATGTATAATGGAGcC |
| cj1429c-MASCR1 | CCCATCTTGCTCCTCAGGATTGC |
| cj1437c-MASCF1 | AGGCTTTTGCATTGGCGAGT |
| cj1437c-MASCmR1M | TTTGCTACAGAATCATTGAATATGCCTCgA |
| cj1437c-MASCmR2M | TTTGCTACAGAATCATTGAATATCCGCgT |

[a]The single-underlined and double-underline nucleotides indicate EcoRI recognition sites and mutated sites, respectively. Lowercase letters in MASC primer sequences indicate mismatched bases that were used to improve the specificity by destabilizing the 3′-end of the non-allelic primer-template complex.
