## Supplementary material for "Stabilizing genetically unstable simple sequence repeats in the *Campylobacter jejuni* genome by multiplex genome editing: a reliable approach for delineating multiple phase-variable genes": S2 Table

**S2 Table. Specific combinations of template DNA and primers used to amplify donor DNA templates**

| First-step PCR | | Second-step PCR | | Generated PCR product (length of homologous region) |
|---|---|---|---|---|
| Template | Primers | Template | Primers | |
| Chromosomal DNA NCTC11168 | cj1339c(flaA)-f1E and cj1339c-cat-r1 | First-step PCR products | cj1339c(flaA)-f1E and cj1339c(flaA)-r2E | Δ*flaA*::*cat*-1 (1 kb) |
| pSYC-*cat* | c-cat-f1 and c-cat-r2 | | | |
| Chromosomal DNA NCTC11168 | cat-cj1339c(flaA)-f2 and cj1339c(flaA)-r2E | | | |
| Chromosomal DNA NCTC11168 | cj1339c(flaA)-f1E and cj1339c-kan-r1 | First-step PCR products | cj1339c(flaA)-f1E and cj1339c(flaA)-r1E | Δ*flaA*::*kan*-1 (1 kb) |
| pSYC-*kan* | c-kan-f1 and c-kan-r1 | | | |
| Chromosomal DNA NCTC11168 | kan-cj1339c(flaA)-f1 and cj1339c(flaA)-r1E | | | |
| Chromosomal DNA NCTC11168 | cj1339c(flaA)-f1E and cj1339c(flaA)-r2E | | | *flaA*$^{+}$-1 (1 kb) |
| Chromosomal DNA NCTC11168 | cj1673c-f1000E and 176_1669-pUCFa-r1 | First-step PCR products | cj1673c-f1000E and cj1673c-r1000E | Δ*recA*::*cat*-1 (1 kb) |
| pSYC-*cat* | pUCFa-f1 and pUCFa-r1 | | | |

| | | | | |
|---|---|---|---|---|
| Chromosomal DNA NCTC11168 | pUCFa-176_1669-f1 and cj1673c-r1000E | | | |
| Chromosomal DNA SYC1003 | 176rpsLmt-f50E and 176rpsLmt-r50E | | | *rpsL*$^{K88R}$-1 (50 b) |
| Chromosomal DNA SYC1003 | 176rpsLmt-f100E and 176rpsLmt-r100E | | | *rpsL*$^{K88R}$-2 (100 b) |
| Chromosomal DNA SYC1003 | 68rpsLmt-f500 and 176rpsLmt-r500 | | | *rpsL*$^{K88R}$-3-1 (500 b) |
| Chromosomal DNA SYC1003 | 68rpsLmt-f500E and 176rpsLmt-r500 | | | *rpsL*$^{K88R}$-4-1 (500 b) |
| Chromosomal DNA SYC1003 | 68rpsLmt-f500E and 176rpsLmt-r500E | | | *rpsL*$^{K88R}$-5-1 (500 b) |
| Chromosomal DNA SYC1003 | 68rpsLmt-f1000 and 176rpsLmt-r1000 | | | *rpsL*$^{K88R}$-6-1 (1 kb) |
| Chromosomal DNA SYC1003 | 68rpsLmt-f1000E and 176rpsLmt-r1000 | | | *rpsL*$^{K88R}$-7-1 (1 kb) |
| Chromosomal DNA NCTC11168 | 68rpsLmt-f1000E and rpsL(CJ0491)-StmR-R2 | First-step PCR products | 68rpsLmt-f1000E and 176rpsLmt-r1000E | *rpsL*$^{K88R}$-8-1 (1 kb) |
| Chromosomal DNA NCTC11168 | rpsL(CJ0491)-StmR-F2 and 176rpsLmt-r1000E | | | |
| Chromosomal DNA SYC1003 | 68rpsLmt-f2000E and 176rpsLmt-r2000E | | | *rpsL*$^{K88R}$-9-1 (2 kb) |
| Chromosomal DNA | 68rpsLmt-f1000E and | | | *rpsL*$^{+}$-8-1 (1 kb) |

| | | | | |
|---|---|---|---|---|
| NCTC11168 | 176rpsLmt-r1000E | | | |
| Chromosomal DNA<br>NCTC11168 | 68rpsLmt-f2000E and<br>176rpsLmt-r2000E | | | *rpsL*$^{+}$-9-1 (2 kb) |
| Chromosomal DNA<br>NCTC11168 | cj1426c-f1E and cj1426c-<br>ON-r1 | First-step PCR<br>products | cj1426c-f1E and<br>cj1426c-r1E | *cj1426*$^{\mathrm{ON}}$ (2 kb) |
| Chromosomal DNA<br>NCTC11168 | cj1426c-ON-f1 and<br>cj1426c-r1E | | | |
| Chromosomal DNA<br>NCTC11168 | cj1426c-f1E and cj1426c-<br>OFF(-1)-r2 | First-step PCR<br>products | cj1426c-f1E and<br>cj1426c-r1E | *cj1426*$^{\mathrm{OFF}}$ (2 kb) |
| Chromosomal DNA<br>NCTC11168 | cj1426c-OFF(-1)-f2 and<br>cj1426c-r1E | | | |
| Chromosomal DNA<br>NCTC11168 | cj1429c-f3E and cj1429c-<br>ON-r2 | First-step PCR<br>products | cj1429c-f3E and<br>cj1429c-r1E | *cj1429*$^{\mathrm{ON}}$ (2 kb) |
| Chromosomal DNA<br>NCTC11168 | cj1429c-ON-f1 and<br>cj1429c-r1E | | | |
| Chromosomal DNA<br>NCTC11168 | cj1429c-f3E and cj1429c-<br>OFF(-1)-r2 | First-step PCR<br>products | cj1429c-f3E and<br>cj1429c-r1E | *cj1429*$^{\mathrm{OFF}}$ (2 kb) |
| Chromosomal DNA<br>NCTC11168 | cj1429c-OFF(-1)-f2 and<br>cj1429c-r1E | | | |
| Chromosomal DNA<br>NCTC11168 | cj1139c-f2E and cj1139c-<br>ON-r1 | First-step PCR<br>products | cj1139c-f2E and<br>cj1139c-r2E | *cj1139*$^{\mathrm{ON}}$ (2 kb) |
| Chromosomal DNA<br>NCTC11168 | cj1139c-ON-f1 and<br>cj1139c-r2E | | | |

| | | | | |
|---|---|---|---|---|
| Chromosomal DNA NCTC11168 | cj1139c-f2E and cj1139c-OFF(-1)-r2 | First-step PCR products | cj1139c-f2E and cj1139c-r2E | *cj1139*$^{\text{OFF}}$ (2 kb) |
| Chromosomal DNA NCTC11168 | cj1139c-OFF(-1)-f2 and cj1139c-r2E | | | |
| Chromosomal DNA NCTC11168 | cj1420c-f1E and cj1420c -ON-r1 | First-step PCR products | cj1420c-f1E and cj1420c-r1E | *cj1420*$^{\text{ON}}$ (2 kb) |
| Chromosomal DNA NCTC11168 | cj1420c -ON-f1 and cj1420c-r1E | | | |
| Chromosomal DNA NCTC11168 | cj1420c-f1E and cj1420c -OFF(-1)-r2 | First-step PCR products | cj1420c-f1E and cj1420c-r1E | *cj1420*$^{\text{OFF}}$ (2 kb) |
| Chromosomal DNA NCTC11168 | cj1420c -OFF(-1)-f2 and cj1420c-r1E | | | |
| Chromosomal DNA NCTC11168 | cj1145c-f1E and cj1145c -ON-r1 | First-step PCR products | cj1145c-f1E and cj1145c-r1E | *cj1144*$^{\text{ON}}$ (2 kb) |
| Chromosomal DNA NCTC11168 | cj1145c -ON-f1 and cj1145c -r1E | | | |
| Chromosomal DNA NCTC11168 | cj1145c-f1E and cj1145c -OFF(-1)-r2 | First-step PCR products | cj1145c-f1E and cj1145c-r1E | *cj1144*$^{\text{OFF}}$ (2 kb) |
| Chromosomal DNA NCTC11168 | cj1145c - OFF(-1)-f2 and cj1145c -r1E | | | |
| Chromosomal DNA NCTC11168 | cj1437c-f1E and cj1437c -ON-r1 | First-step PCR products | cj1437c-f1E and cj1437c-r1E | *cj1437*$^{\text{ON}}$ (2 kb) |
| Chromosomal DNA | cj1437c -ON-f1 and | | | |

| | | | | |
|---|---|---|---|---|
| NCTC11168 | cj1437c -r1E | | | |
| Chromosomal DNA NCTC11168 | cj1437c-f1E and cj1437c -OFF(-1)-r3 | First-step PCR products | cj1437c-f1E and cj1437c-r1E | *cj1437*$^{\mathrm{OFF}}$ (2 kb) |
| Chromosomal DNA NCTC11168 | cj1437c -OFF(-1)-f3 and cj1437c -r1E | | | |
| Chromosomal DNA NCTC11168 | cj1422c-f1E and cj1421/22cc -ON-r1 | First-step PCR products | cj1422c-f1E and cj1422c-r2E | *cj1422*$^{\mathrm{ON}}$ (2 kb) |
| Chromosomal DNA NCTC11168 | cj1421/22cc -ON-f1 and cj1422c-r2E | | | |
| Chromosomal DNA NCTC11168 | cj1422c-f1E and cj1421/22cc -OFF(-1)-r2 | First-step PCR products | cj1422c-f1E and cj1422c-r2E | *cj1422*$^{\mathrm{OFF}}$ (2 kb) |
| Chromosomal DNA NCTC11168 | cj1421/22cc -OFF(-1)-f2 and cj1422c-r2E | | | |
| Chromosomal DNA NCTC11168 | cj1422c-f2E and cj1421/22cc -ON-r1 | First-step PCR products | cj1422c-f2E and cj1422c-r1E | *cj1421*$^{\mathrm{ON}}$ (2 kb) |
| Chromosomal DNA NCTC11168 | cj1421/22cc -ON-f1 and cj1422c-r1E | | | |
| Chromosomal DNA NCTC11168 | cj1422c-f2E and cj1421/22cc -OFF(-1)-r2 | First-step PCR products | cj1422c-f2E and cj1422c-r1E | *cj1421*$^{\mathrm{OFF}}$ (2 kb) |
| Chromosomal DNA NCTC11168 | cj1421/22cc -OFF(-1)-f2 and cj1422c-r1E | | | |
| Chromosomal DNA 81-176 | cjj81176_1339-f1E and flaA81176-cat-r1 | First-step PCR products | cjj81176_1339-f1E and cjj81176_1339-r1E | Δ*flaA*::*cat*-2(1 kb) |

| | | | | |
|---|---|---|---|---|
| pSYC-*cat* | c-cat-f1 and c-cat-r2 | | | |
| Chromosomal DNA 81-176 | cat-flaA81176-f2 and cjj81176_1339-r1E | | | |
| Chromosomal DNA 81-176 | cjj81176_1339-f1E and flaA81176-kan-f1 | First-step PCR products | cjj81176_1339-f1E and cjj81176_1339-r1E | Δ*flaA*::*kan*-2(1 kb) |
| pSYC-*kan* | c-kan-f1 and c-kan-r1 | | | |
| Chromosomal DNA 81-176 | kan-flaA81176-f1 and cjj81176_1339-r1E | | | |
| Chromosomal DNA 81-176 | 176_1669-f1000E and 176_1669-pUCFa-r1 | First-step PCR products | 176_1669-f1000E and 176_1669-r1000E | Δ*recA*::*cat*-2(1 kb) |
| pSYC-*cat* | pUCFa-f1 and pUCFa-r1 | | | |
| Chromosomal DNA 81-176 | pUCFa-176_1669-f1 and 176_1669-r1000E | | | |
| Chromosomal DNA SYC2003 | 176rpsLmt-f500 and 176rpsLmt-r500 | | | *rpsL*$^{K88R}$-3-2 (500 b) |
| Chromosomal DNA SYC2003 | 176rpsLmt-f500E and 176rpsLmt-r500 | | | *rpsL*$^{K88R}$-4-2 (500 b) |
| Chromosomal DNA SYC2003 | 176rpsLmt-f500E and 176rpsLmt-r500E | | | *rpsL*$^{K88R}$-5-2 (500 b) |
| Chromosomal DNA SYC2003 | 176rpsLmt-f1000 and 176rpsLmt-r1000 | | | *rpsL*$^{K88R}$-6-2 (1 kb) |
| Chromosomal DNA SYC2003 | 176rpsLmt-f1000E and 176rpsLmt-r1000 | | | *rpsL*$^{K88R}$-7-2 (1 kb) |

| | | | | |
|---|---|---|---|---|
| Chromosomal DNA 81-176 | 176rpsLmt-f1000E and rpsL(CJ0491)-StmR-R2 | First-step PCR products | 176rpsLmt-f1000E and 176rpsLmt-r1000E | *rpsL*$^{K88R}$-8-2 (1 kb) |
| Chromosomal DNA 81-176 | rpsL(CJ0491)-StmR-F2 and 176rpsLmt-r1000E | | | |
| Chromosomal DNA SYC2003 | 176rpsLmt-f2000E and 176rpsLmt-r2000E | | | *rpsL*$^{K88R}$-9-2 (1 kb) |
| Chromosomal DNA 81-176 | 176rpsLmt-f1000E and 176rpsLmt-r1000E | | | *rpsL*$^{+}$-8-2 (1 kb) |
| Chromosomal DNA 81-176 | 176rpsLmt-f2000E and 176rpsLmt-r2000E | | | *rpsL*$^{+}$-9-2 (2 kb) |
| Chromosomal DNA SYC1P255 | cj1426c-f1E and cj1426c-astA-r2 | First-step PCR products | cj1426c-f1E and cj1426c-r1E | *cj1426*$^{ON}$::*astA* (2 kb) |
| Chromosomal DNA 81-176 | astA-f2 and astA-r2 | | | |
| Chromosomal DNA NCTC11168 | astA-cj1426c-f2 and cj1426c-r1E | | | |
| Chromosomal DNA SYC1007 | cj1426c-f1E and cj1426c-WT-r1 | First-step PCR products | cj1426c-f1E and cj1426c-r1E | *cj1426*::*astA* (2 kb) |
| Chromosomal DNA SYC1007 | cj1426c-WT-f1 and cj1426c-r1E | | | |
| Chromosomal DNA SYC1007 | cj1426c-f1E and cj1426c-OFF(-1)-r2 | First-step PCR products | cj1426c-f1E and cj1426c-r1E | *cj1426*$^{OFF}$::*astA* (2 kb) |
| Chromosomal DNA | cj1426c-OFF(-1)-f2 and | | | |

| | |
|---|---|
| SYC1007 | cj1426c-r1E |
