## Supplementary material for "Stabilizing genetically unstable simple sequence repeats in the *Campylobacter jejuni* genome by multiplex genome editing: a reliable approach for delineating multiple phase-variable genes": S3 Table

**S3 Table. Specific combinations of donor DNA molecules and recipient strains used for natural transformation**

| Donor DNA (PCR fragment) | Recipient strain | Resulting strain |
|---|---|---|
| Δ*flaA*::*cat*-1 | NCTC11168 | SYC1001 |
| Δ*flaA*::*kan*-1 | NCTC11168 | SYC1002 |
| *rpsL*$^{K88R}$-8-1 | NCTC11168 | SYC1003 |
| Δ*recA*::*cat*-1 | NCTC11168 | SYC1004 |
| *cj1426*::*astA*, Δ*flaA*::*kan*-1 | SYC1007 | SYC1006 |
| *cj1426*$^{ON}$::*astA*, Δ*flaA*::*cat*-1 | NCTC11168 | SYC1007 |
| *cj1426*$^{OFF}$::*astA*, Δ*flaA*::*kan*-1 | SYC1007 | SYC1008 |
| Δ*flaA*::*kan*-1 (or Δ*flaA*::*cat*-1), *cj1139*$^{OFF}$, *cj1144*$^{OFF}$, *cj1420*$^{OFF}$, *cj1421*$^{OFF}$, *cj1422*$^{OFF}$, *cj1426*$^{OFF}$, *cj1429*$^{OFF}$, *cj1437*$^{OFF}$ | NCTC11168 | SYC1P000K |
| *flaA*$^{+}$-1 | SYC1P000K | SYC1P000 |
| Δ*flaA*::*cat*-1, *cj1420*$^{OFF}$, *cj1426*$^{OFF}$ | SYC1P037 | SYC1P001C |
| Δ*flaA*::*kan*-1, *cj1420*$^{OFF}$ | SYC1P036C | SYC1P004K |
| Δ*flaA*::*cat*-1, *cj1420*$^{OFF}$ | SYC1P037 | SYC1P005C |
| Δ*flaA*::*kan*-1, *cj1426*$^{OFF}$ | SYC1P036C | SYC1P032K |
| Δ*flaA*::*cat*-1, *cj1426*$^{OFF}$ | SYC1P037 | SYC1P033C |
| Δ*flaA*::*cat*-1, *cj1437*$^{OFF}$ | SYC1P037 | SYC1P036C |
| Δ*flaA*::*cat*-1, *cj1139*$^{OFF}$, *cj1144*$^{OFF}$, *cj1421*$^{OFF}$, *cj1422*$^{OFF}$, *cj1429*$^{OFF}$ | SYC1P255 | SYC1P037C |
| *flaA*$^{+}$-1 | SYC1P037C | SYC1P037 |
| Δ*flaA*::*kan*-1 (or Δ*flaA*::*cat*-1), *cj1139*$^{ON}$, *cj1144*$^{ON}$, *cj1420*$^{ON}$, *cj1421*$^{ON}$, *cj1422*$^{ON}$, *cj1426*$^{ON}$, *cj1429*$^{ON}$, *cj1437*$^{ON}$ | NCTC11168 | SYC1P255K |
| *flaA*$^{+}$-1 | SYC1P255K | SYC1P255 |
| Δ*flaA*::*cat*-2 | 81-176 | SYC2001 |
| Δ*flaA*::*kan*-2 | 81-176 | SYC2002 |
| *rpsL*$^{K88R}$-8-2 | 81-176 | SYC2003 |
| Δ*recA*::*cat*-2 | 81-176 | SYC2004 |
