## Supplementary material for "Stabilizing genetically unstable simple sequence repeats in the *Campylobacter jejuni* genome by multiplex genome editing: a reliable approach for delineating multiple phase-variable genes": S4 Table

**S4 Table. Primer mixes used for MASC PCR**

| Name | Size (bp) | ON/OFF detected | Phase |
|---|---|---|---|
| Mix ON1 | | | |
| cj1422c-MASCF4 and cj1421c/22c-MASCmR1M | 449 | *cj1422*$^{ON}$ | |
| cj1421c-MASCF1 and cj1421c/22c-MASCmR1M | 319 | *cj1421*$^{ON}$ | |
| cj1437c-MASCF1 and cj1437c-MASCmR1M | 250 | *cj1437*$^{ON}$ | |
| cj1145c-MASCmF1M and cj1145c-MASCR1 | 150 | *cj1144*$^{ON}$ | |
| Mix ON2 | | | |
| cj1429c-MASCmF1M and cj1429c-MASCR1 | 400 | *cj1429*$^{ON}$ | |
| cj1426c-MASCmF1M and cj1426c-MASCR1 | 300 | *cj1426*$^{ON}$ | |
| cj1139c-MASCmF2M and cj1139c-MASCR2 | 200 | *cj1139*$^{ON}$ | |
| cj1420c-MASCmF1M and cj1420c-MASCR1 | 100 | *cj1420*$^{ON}$ | |
| Mix OFF1 | | | |
| cj1422c-MASCF4 and cj1421c/22c-MASCmR2M | 449 | *cj1422*$^{OFF}$ | |
| cj1421c-MASCF1 and cj1421c/22c-MASCmR2M | 319 | *cj1421*$^{OFF}$ | |
| cj1437c-MASCF1 and cj1437c-MASCmR2M | 250 | *cj1437*$^{OFF}$ | |
| cj1145c-MASCmF2M and cj1145c-MASCR1 | 150 | *cj1144*$^{OFF}$ | |
| Mix OFF2 | | | |
| cj1429c-MASCmF2M and cj1426c-MASCR1 | 400 | *cj1429*$^{OFF}$ | |

| | | |
|---|---|---|
| cj1426c-MASCmF2M and cj1426c-MASCR1 | 300 | *cj1426*$^{OFF}$ |
| cj1139c-MASCmF3M and cj1139c-MASCR2 | 200 | *cj1139*$^{OFF}$ |
| cj1420c-MASCmF2M and cj1420c-MASCR1 | 100 | *cj1420*$^{OFF}$ |
