## Supplementary material for "Stabilizing genetically unstable simple sequence repeats in the *Campylobacter jejuni* genome by multiplex genome editing: a reliable approach for delineating multiple phase-variable genes": S5 Table

**S5 Table. Specific combinations of target genes and primers in PPT-Seq**

| Target gene | PCR primers | Sequencing primer |
|---|---|---|
| *cj1139* | cj1139c-f2E and cj1139c-r2E | cj1139c-MASCR2 |
| *cj1144* | cj1145c-f1E and cj1145c-r1E | cj1145c-MASCR1 |
| *cj1420* | cj1420c-f1E and cj1420c-r1E | cj1420c-MASCR1 |
| *cj1421* | cj1422c-f2E and cj1422c-r1E | cj1421c-MASCF1 |
| *cj1422* | cj1422c-f1E and cj1422c-r2E | cj1422c-MASCF4 |
| *cj1426* | cj1426c-f1E and cj1426c-r1E | cj1426c-MASCR1 |
| *cj1429* | cj1429c-f3E and cj1429c-r1E | cj1429c-MASCR1 |
| *cj1437* | cj1437c-f1E and cj1437c-r1E | cj1437c-MASCF1 |
