## Supplementary material for "Stabilizing genetically unstable simple sequence repeats in the *Campylobacter jejuni* genome by multiplex genome editing: a reliable approach for delineating multiple phase-variable genes": S6 table

**S6 Table. Primer sets used for allele-specific PCR and sequencing of *cj1426*::*astA* translational fusions**

| Reporter gene | Allele-specific PCR primers | Sequencing | |
|---|---|---|---|
| | | PCR primers | Sequencing primer |
| *cj1426*::*astA* | cj1426c-MASCwF1M and astA-MASCR1 | cj1426c-f1E and astA-MASCR1 | astA-MASCR1 |
| *cj1426*$_{\mathrm{ON}}$::*astA* | cj1426c-MASCmF1M and astA-MASCR1 | cj1426c-f1E and astA-MASCR1 | astA-MASCR1 |
| *cj1426*$_{\mathrm{OFF}}$::*astA* | cj1426c-MASCmF2M and astA-MASCR1 | cj1426c-f1E and astA-MASCR1 | astA-MASCR1 |
